# Oligosaccharyltransferase (OST) complex inhibition effectively treats rodent and human prions

**DOI:** 10.64898/2025.12.07.692846

**Authors:** Kathryn S. Beauchemin, Judit Kun, Bradley Groveman, Katie Williams, Francesca Salerno, Lisa M. Francomacaro, Patrick Robison, Cathryn L. Haigh, Surachai Supattapone

**Affiliations:** Departments of Biochemistry, Geisel School of Medicine at Dartmouth, Hanover, New Hampshire 03755, USA; Medicine, Geisel School of Medicine at Dartmouth, Hanover, New Hampshire 03755, USA; Prion Cell Biology Section, Laboratory of Neurological Infections and Immunity, Rocky Mountain Laboratories, National Institute of Allergy and Infectious Diseases, National Institutes of Health, Hamilton, Montana, United States of America

**Keywords:** N-glycosylation, sCJD, target, therapy, PrP^C^

## Abstract

Prion diseases are invariably fatal neurodegenerative diseases that occur when the prion protein misfolds into a pathogenic form. There are currently no clinical treatments or cures for prion disease. Current challenges in the development of prion therapeutics include prion strain specificity, which can cause the emergence of drug-resistant prions, and lack of efficacy in treating human prions despite promising results in rodent models. Here we identify a novel therapeutic target for prion disease: the oligosaccharyltransferase (OST) complex. The OST complex is responsible for transferring the mature glycan to the acceptor polypeptide during Nglycosylation. We found that inhibiting OST effectively treats rodent prions in various dividing and non-dividing cell types. Importantly, we also demonstrate efficacy in treating human sCJD prions in non-dividing cerebral organoids. Inhibition of OST results in a 50% reduction in cell surface expression of the prion protein, PrP^C^. In addition, lysates of cells treated with the OST inhibitor NGI-1 were unable to amplify PrP^Sc^ seeds in Protein Misfolding Cyclic Amplification (PMCA) reactions. In summary, our results identify OST as a novel therapeutic target that regulates both the abundance of cell surface PrP^C^ as well as its ability to convert into multiple strains of PrP^Sc^, including human prions, in various *in vitro* systems.

**Author Summary:** Prion diseases, such as Creutzfeldt–Jakob disease, are fatal brain disorders caused when a normal protein (PrP^C^) misfolds into a harmful form that spreads through the brain. There are no effective treatments, and drug development has been hampered by “strain” differences in prions that can lead to resistance and by therapies that work in rodents but not in humans. This study identifies a new treatment target: the oligosaccharyltransferase (OST) complex, a cellular machine that adds sugar groups to proteins. Blocking OST with a small molecule (NGI1) limited prion growth in multiple types of rodent cells, including non-dividing cells, and— critically—also worked against human sporadic CJD prions in laboratory-grown human brain organoids. OST inhibition cut the amount of normal prion protein on cell surfaces in half, reducing the raw material available to convert into the disease-causing form. In addition, extracts from NGI-1–treated cells could no longer drive prion formation in a sensitive lab amplification test.

These results suggest that targeting OST may offer a new strategy that works across different prion strains, including human ones.

## Introduction

Prion diseases are transmissible, incurable, invariably fatal neurodegenerative diseases that occur in humans and other mammals[1]. Prion diseases occur when the host-encoded prion protein (PrP^C^) misfolds into a pathogenic form (PrP^Sc^), which can template further PrP^C^ misfolding[2].

PrP^C^ is 30-40kDa glycoprotein which is attached to the outer leaflet of the plasma membrane by a glycophosphatidylinositol (GPI) anchor[3]. PrP^C^ has two sites for asparagine-linked (N-linked) glycosylation and exists as un-glycosylated, mono-glycosylated, or di-glycosylated glycoprotein of complex type in equal proportion in the brain[4]. The N-glycans on PrP^C^ are branched and terminally sialylated[5–8]. Although much effort has gone into understanding the impact of Nglycosylation on the expression and localization of PrP^C^, work *in vivo* or in cultured cells has been technically limited due to the requirement for genetic manipulation of PrP^C^. Numerous studies in which the AsnXaaThr or AsnXaaSer sites of PrP^C^ are genetically modified have resulted in conflicting or inconsistent results, which may be attributable to differences in the cell type/model used, species-specific differences in PrP^C^ sequence, and amino acid substitutions used[9–14].

Prion strains are different misfolded conformations of PrP^Sc^ which are distinguishable biochemically, symptomatically, and by strain-specific neurotropism[15, 16]. The ratio of un-, mono-, and di-glycosylated proteinase K (PK)-resistant PrP^Sc^ is characteristic of a given prion strain and can be used to help distinguish certain prion strains by western blotting[17]. Additionally, prion strains have different glycosylation preferences[18, 19]. For example, mouse prion strains require un-glycosylated PrP^C^ substrate for prion conversion *in vitro*, whereas hamster strains require di-glycosylated PrP^C^ substrate.

Decades of work have gone into the development of therapeutics for prion disease, yet, to date, there are no efficacious treatments or prophylactics for any type of prion disease (reviewed in [20]). Significant challenges in prion therapy include strain dependency of anti-prion therapeutics[21–26] which can cause the emergence of drug-resistant prions[22, 23, 27–30], reduction in efficacy for established infections[22, 25, 31–33], and ineffectiveness in treating human prions despite promising results in rodent models of disease[20, 24–26, 34, 35]. An intuitive and attractive strategy for the development of prion therapeutics involves limiting the availability of substrate PrP^C^. Since PrP^C^ is required for prion conversion[36], the challenges of prion strain specificity, species specificity, and drug resistance would likely be avoided using this approach. Additionally, PrP^C^-limiting therapies may be effective in treating all forms of prion disease (sporadic, familial, and acquired). Recent successes in the development of such therapies using Zinc Finger Repressors[37], antisense oligonucleotides[33, 38, 39], liposomesiRNA-peptide complexes[40], divalent siRNA[41], *in vivo* base editing[42], and epigenetic modifiers[43] underscore the practicality of this approach and provide great hope for the future of prion therapeutics. Additionally, PrP^C^ must be present at the cell surface for *de novo* prion infection to occur, to maintain chronic prion infection, and to induce neurotoxicity[36, 44–46], suggesting that therapeutic depletion of PrP^C^ by interfering with its normal metabolic processes may be a viable alternative approach to direct genetic manipulation of PrP^C^.

We recently performed a genome-wide CRISPR/Cas9 knockout screen in the prion-infectible mouse neuronal-like cell line, CAD5, and identified the N-glycosylation pathway as an important positive regulator of PrP^C^ cell surface expression. One critical step in the N-glycosylation biosynthetic pathway involves the transfer of the mature glycan (Glc_₃_Man_₉_GlcNAc_₂_) to the acceptor polypeptide in the endoplasmic reticulum, which is performed by the enzyme complex oligosaccharyltransferase (OST)[47–49]. OST exists as two isoforms, STT3A-OST and STT3BOST based on the catalytic subunit used (STT3A or STT3B). Since PrP^C^ glycosylation is dependent on the glycan transfer step mediated by OST and this occurs further upstream in the N-glycosylation pathway than the enzymatic steps of other N-glycosylation inhibitor molecules previously tested in the context of PrP^C^ or PrP^Sc^[8, 50], we were interested to test the effect of NGI-1, a recently identified potent small molecule inhibitor of STT3A and STT3B[51–53], on PrP^C^ surface expression and PrP^Sc^ levels in cultured cells and organoids.

## Materials and Methods

### Cell lines and cell culture

WT CAD5 cells were kindly provided by Charles Weissmann (Scripps Florida, Jupiter, FL, USA). CAD5 cells genetically engineered using CRISPR/Cas9 to lack endogenous mouse PrP expression (CAD5 PrP KO) and CAD5 PrP KO cells engineered to stably express hamster PrP (CAD5 HaPrP) were kindly provided by Joel Watts (University of Toronto, Toronto, Canada). Rabbit kidney epithelial RK13 cells expressing mouse PrP under the Tet-On system (moRK13) were kindly provided by Didier Vilette (University of Toulouse, Toulouse, France).

CAD5 cells were maintained in Opti-MEM I Reduced-Serum Media with GlutaMAX (Gibco, Waltham, MA, USA) supplemented with 10% HyClone Bovine Growth Serum (BGS) (Cytiva, Marlborough, MA, USA) and 1X penicillin/streptomycin (Corning, Corning, NY, USA). Prion-infected CAD5 cells were cultured using 0.2X penicillin/streptomycin. To differentiate CAD5 cells, cells were rinsed with PBS and complete media was replaced with DMEM:F12 (Gibco) supplemented with 50 ng/mL sodium selenite (Sigma-Aldrich, St. Louis, MO, USA). Differentiation was considered complete after 4 days incubation in the differentiation media. BE(2)-C human neuroblastoma cells (ATCC CRL-2268) were cultured in a 1:1 mix of EMEM (ATCC) and F-12 (Gibco) media supplemented with 10% fetal bovine serum (Cytiva) at 37°C with 5% CO_2_ until >90% confluent then split 1:4-1:10 with 0.05% Trypsin / 0.53 mM EDTA (Corning). BE(2)-C differentiation was performed by adding all-trans retinoic acid (Thermo Fisher Scientific, Waltham, MA, USA) to a final concentration of 10 μM in complete growth media and was considered complete after 4 days of incubation.

WTC11-NGN2 induced pluripotent stem cells (iPSCs) [54] were a gift of Li Gan (Weill Cornell, New York, NY, USA) and were cultured as recommended by Fernandopulle et al.[55]. Plates were coated with Matrigel (Corning) as in the established protocol. Cells were cultured in complete mTeSR1 medium. To make complete mTeSR1 media, mTeSR1 5X media supplement (STEMCELL Technologies, Vancouver, Canada) was warmed to room temperature and added to mTeSR1 basal medium (STEMCELL Technologies). Aliquots of complete mTeSR1 media were stored at -20 °C and used within two weeks after thawing with storage at 4 °C. iPSCs were thawed from liquid nitrogen storage at 37 °C and complete mTeSR1 media was added dropwise. Cells were centrifuged at 240 *x g* for 5 min at room temperature and resuspended gently in complete mTeSR1 with 10 μM ROCK inhibitor (STEMCELL Technologies). After 24 hr incubation at 37 °C, 5% CO_2_, media was replaced with fresh complete mTeSR1 without ROCK inhibitor. Culture medium was changed daily until reaching 80% confluence. To split cells, cells were washed with PBS and incubated for 30 seconds at room temperature with ReLeSR (STEMCELL Technologies). ReLeSR was removed and dry cells were incubated at 37 °C for 7 min before cells were dislodged gently with complete media. Cells were split 1:4 to 1:6 with addition of complete media.

WTC11-NGN2 iPSCs (kindly provided by Li Gan, Gladstone Institute, San Francisco, CA through the Tau Consortium) were induced to differentiate into neurons over the course of five days (numbered 0 to 4, referring to days post-differentiation induction). On day zero, undifferentiated iPSCs were rinsed with PBS, lifted using Accutase (STEMCELL Technologies) plus 1X DNase, and centrifuged at 200 *x g* for 5 min at room temperature. The cell pellet was resuspended in induction media containing 2 μg/mL doxycycline and 10 μM ROCK inhibitor and incubated at 37°C, 5% CO_2_ for 24 hr. Induction media consisted of DMEM/F12, HEPES (Gibco) with 1X N2 supplement (Gibco), 1X Non-essential Amino Acids (Gibco), and1X L-glutamine (Gibco). On day one, cells were washed with PBS and switched to induction media plus 2 μg/mL doxycycline (no ROCK inhibitor). On day two, cells were washed 2X with PBS and refreshed with induction media plus 2 μg/mL doxycycline (no ROCK inhibitor). On day three, cells were lifted using Accutase for 3 min at room temperature. Cells were centrifuged at 200 *x g* for 5 min at room temperature, resuspended in freezing medium (90% knockout serum, 10% DMSO), and stored at -80 °C for 24 hr before transferring to liquid nitrogen storage.

Day 3 differentiated neurons (i^3^Neurons) were cultured as previously described on poly-Lornithine (Millipore Sigma) and laminin (Gibco) double-coated plates in cortical neuron culture media consisting of BrainPhys neuronal medium (Stem Cell Technologies), B27 supplement (Gibco), 10 ng/mL BDNF (PeproTech, Cranbury, NJ, USA), 10 ng/mL NT-3 (PeproTech), and 1 μg/mL laminin (Gibco)[55]. Cells underwent half-medium changes every 3-4 days with dropwise addition until day 14, at which point neurons were mature glutamatergic cortical neurons.

moRK13 cells were maintained in Opti-MEM I Reduced-Serum Media with GlutaMAX (Gibco) supplemented with 10% fetal bovine serum (Cytiva) and 1X penicillin/streptomycin (Corning). Prion-infected RK13 cells were cultured using 0.2X penicillin/streptomycin. To induce expression of PrP, doxycycline (Sigma-Aldrich) was added to a final concentration of 1 µg/mL in complete growth media. Complete growth media containing doxycycline was refreshed every 48-72 hr to ensure continuous expression of PrP.

Cell lines were routinely monitored for mycoplasma contamination using the LookOut Mycoplasma PCR Detection Kit (Sigma-Aldrich, St. Louis, MO, USA).

### N-glycosylation inhibitor treatments

Unless otherwise noted, all experiments involving cell treatment with N-glycosylation inhibitors underwent treatment at the indicated final concentration in complete cell culture media, with media/drug refreshes every 48-72 hr.

Kifunensine (Sigma-Aldrich, K1140) and NGI-1 (Sigma-Aldrich, SML1620) were prepared as 10 mM stock solutions, kifunensine in cell culture grade water (Corning) and NGI-1 in DMSO (Sigma-Aldrich, D2650). Both drugs were aliquoted and frozen at manufacturer’s recommended storage temperature. Thawed aliquots were discarded after use to avoid repeated freeze/thaw.

### Flow cytometry for surface PrP^C^

All flow cytometry experiments were performed on a CytoFLEX flow cytometer (Beckman Coulter, Brea, CA, USA).

To detect surface PrP^C^ in CAD5 cells, the anti-PrP antibody 6D11 (Biolegend, San Diego, CA, USA) was used at 0.8 µg per sample. The anti-PrP antibody 4D5 (Invitrogen) was used at a 1:200 dilution to detect surface PrP^C^ in the i^3^Neurons. Each sample contained approximately 500,000 cells in 100 µL PBS/2% FBS (staining buffer). Primary incubations were performed at 4 °C for 1 hr, followed by three washes in PBS, with final resuspension to 100 µL in staining buffer.

Anti-Mouse IgG2a Secondary Antibody:PE (Invitrogen, 12-4210-82) was used as the secondary antibody at the manufacturer’s recommended dilution for CAD5 staining. Rat anti-mouse IgG1:APC secondary antibody (Invitrogen) was used for i^3^N staining. Secondary incubations were performed at 4 °C for 30 min, followed by three washes in PBS, with final resuspension in 250 μL staining buffer.

FlowJo software (BD Biosciences, Franklin Lakes, NJ, USA) was used for all flow cytometry analysis. Sequential gating using FSC and SSC characteristics were performed to identify single cell populations. Median fluorescence intensities of the resulting single cell populations were used for comparisons between populations and treatments.

### Ethics statement

The Guide for the Care and Use of Laboratory Animals of the National Research Council was strictly followed for all animal experiments. All experiments involving mice in this study were conducted in accordance with protocol supa.su.1 as reviewed and approved by Dartmouth College’s Institutional Animal Care and Use Committee, operating under the regulations/guidelines of the NIH Office of Laboratory Animal Welfare (assurance number A3259-01) and the United States Department of Agriculture.

### Prion infection of cultured cells

WT CAD5 cells were infected with mouse prion strains 22L and RML via incubation with brain homogenate from terminally ill animals. 10% (w/v) brain homogenate was prepared by weighing brains of terminally ill mice infected with either the 22L or RML strains and homogenizing for 30 sec in sterile 1X PBS (Corning) using a 4-Place Mini Bead Mill Homogenizer (VWR, Radnor, PA, USA) and sterile 1.4 mm ceramic beads (VWR,10158-552). 10% brain homogenate was stored at -80°C until use. WT CAD5 cells were seeded at a density of 1 x 10^5^ cells per well in a 12-well tissue culture plate (Corning) and incubated at 37°C, 5% CO_2_ overnight. 10% brain homogenate was thawed on ice, centrifuged at 400 x *g* for 30 sec at 4 °C twice to pellet debris, then diluted to 0.5% (w/v) brain homogenate in complete cell culture medium. Cell culture medium was replaced with the medium containing 0.5% (w/v) brain homogenate. Cells were exposed to the inoculum for 72 hr, then passaged at a 1:4 or 1:5 dilution every 2-3 days. Cells were passaged for one month prior to expansion, freezing, and analysis of prion infection status.

CAD5 HaPrP cells stably infected with Hyper or 263K were generated as described by Bourkas et al.[56] and kindly provided by Joel Watts (University of Toronto, Toronto, Canada). To maintain a high level of PrP^Sc^, CAD5 HaPrP Hyper or 263K cells were passaged normally for one month prior to expansion and freezing.

moRK13 cells were infected with mouse prion strain 22L as described for the CAD5 cells above with the following alterations to protocol: MoRK13 cells were seeded in 6-well plates and allowed to grow to complete confluence in the presence of 1 μg/mL doxycycline before addition of the inoculum. Cells were exposed to inoculum for one week, with direct doxycycline addition into inoculum-containing media every 48-72 hrs. Cells were then washed twice with PBS and complete cell culture media containing doxycycline was used for cell maintenance. Chronically infected MoRK13-22L cells used for subsequent experiments were made by splitting these cells 1:6 and allowing cells to grow to complete confluence before each split for three weeks in doxycycline-containing media with regular refreshes every 48-72 hrs. Cells were frozen down in FBS/10% DMSO at -80 °C for 24 hr and then transferred to liquid nitrogen for long-term storage.

### Cell lysis and immunoblotting

To immunoblot for PrP^C^, cells were washed 2X with PBS, lysed in lysis buffer (50 mM Tris, pH 8.0, 150 mM NaCl, 0.5% (w/v) sodium deoxycholate, and 0.5% (v/v) Nonidet P-40), collected, and centrifuged at 250 x *g* for 30 sec to pellet DNA. Lysate supernatant was transferred to fresh tubes. Protein concentrations were determined using a Pierce BCA Protein Assay Kit (Thermo Fisher Scientific) and 30-50 µg total protein was loaded per lane.

To check prion infection status, confluent cells were lysed as described above. After BCA analysis, lysates were digested with 20 µg/mL Proteinase K (PK) (Roche, Basel, Switzerland) for 1 h at 37 °C with 350 rpm shaking. Digestions were stopped by the addition of phenylmethylsulfonyl fluoride (PMSF) (Sigma-Aldrich) to a final concentration of 2 mM. Samples were then ultracentrifuged at 100,000 x *g* for 1 h at 4 °C in a Sorvall Discovery M120 SE Micro-Ultracentrifuge with an S45-A rotor (Thermo Fisher Scientific) in safe-lock tubes (Eppendorf, Hamburg, Germany). Pellets were resuspended in 60 µL Milli-Q water plus 20 μL of 6X Laemmli SDS sample buffer (Bioland Scientific LLC, Paramount, CA, USA) containing 9% (v/v) beta-mercaptoethanol (BME) (Gibco) and boiled for 15 min at 95 °C. All PrP^Sc^ samples were normalized such that the same amount of total protein (500-1000 µg total protein predigest) was loaded between samples on each blot.

SDS-polyacrylamide gel electrophoresis (PAGE) was performed using 1.5 mm 12% polyacrylamide gels with an acrylamide/bisacrylamide ratio of 29:1. The gel was transferred to a methanol-charged polyvinylidene difluoride membrane (Millipore Sigma, Burlington, MA, USA) using a Transblot SD semidry transfer cell (Bio-Rad Laboratories, Hercules, CA, USA). The transfer was set at 2.5 mA/cm^2^ for 45 min. To visualize PrP signal, the membrane was blocked in 5% (w/v) nonfat dry milk (Nestlé, Vevey, Switzerland) in TBST (10 mM Tris, pH 7.1, 150 mM NaCl, 0.1% Tween 20) for 1 h at 4 °C. The blocked membrane was then incubated overnight at 4 °C with anti-PrP 6D11 primary antibody, washed three times for 10 min in TBST, then incubated for 1 h with horseradish peroxidase-labeled sheep-anti-mouse secondary antibody (Cytiva). Membrane was then washed four additional times for 10 min each in TBST. Blots were developed with SuperSignal West Femto (Thermo Fisher Scientific) chemiluminescence substrate, and images were captured digitally using an Azure 600 (Azure Biosystems, Dublin, CA, USA) imaging system. Relative molecular masses were determined by comparison to PageRuler Plus Prestained Protein Ladder (Thermo Fisher Scientific). Digitally captured images were subjected to densitometric analysis using the program ImageJ (U.S. National Institutes of Health, Bethesda, Maryland, USA). Biological replicate measurements were obtained (n=3) and statistical significance was determined using unpaired t-tests using GraphPad Prism version 10.2.1 for Windows (GraphPad Software, Boston, MA, USA).

### PMCA with brain homogenate substrate

Mouse brains were harvested from animals perfused with PBS plus 5 mM EDTA. A 10% (w/v) perfused BH substrate was prepared in PMCA conversion buffer consisting of 1X PBS, 1% (v/v) Triton X-100, 5 mM EDTA, and cOmplete Mini Protease Inhibitors (Roche, Basel, Switzerland). 100 µL samples were made of 90 µL mouse BH substrate plus 1 µL RML seed from cell lysate (below) plus 7 µL PMCA conversion buffer plus 2 µL 500 µM NGI-1 (in DMSO) or DMSO. Master mix was made to minimize variability due to pipetting. Unsonicated samples were immediately frozen at -80 °C. Sonicated samples underwent sonication with 20 sec pulses every 30 min for 24 hr at 37 °C.

RML cell lysate seed for PMCA reactions was made from the lysate of CAD5 cells chronically infected with RML. RML-CAD5 cells were grown in 100 mm-diameter culture plates to 100% confluence, then washed twice with 10 mL PBS. The cells from each plate were harvested in 1mL PBS using a cell scraper, combined, and centrifuged at 150 *x g* for 5 min at 4 °C. The cell pellet was washed with 10 mL cold PBS and centrifuged again before being resuspended in 1 mL cold PMCA conversion buffer. Cell lysate was made by serially passaging through hypodermic needles from 18 to 23 gauges. Lysate was centrifuged at 2,000 *x g* for 5 min at 4 °C and supernatant retained as the RML cell lysate seed for PMCA.

### Cell Lysate PMCA

moRK13 cells were grown to confluence on 150 mm-diameter culture plates to 100% confluence in complete moRK13 media [Opti-MEM I Reduced-Serum Media with GlutaMAX, 10% FBS, 1X penicillin/streptomycin]. To obtain cell lysate from cells expressing PrP, the complete moRK13 media contained 1 μg/mL doxycycline. To obtain cell lysate from cells without PrP expression, moRK13 media did not contain doxycycline. Cells were treated with either 5 μM NGI-1 or DMSO in the media for a total of 96 hr with a media refresh at 48 hr. After treatment, plates were washed with 1X PBS (Corning), dissociated using enzyme-free dissociation buffer (Sigma-Aldrich) per plate with manual cell scraping, and pooled in 50 mL centrifuge tubes. Cells were centrifuged at 150 *x g* for 5 min at 4 °C, resuspended in 50 mL 1X PBS, and centrifuged again. Cell pellets were resuspended in PMCA conversion buffer. Cell lysate substrate was made by serially passaging through hypodermic needles from 18 to 23 gauges. Lysate was centrifuged at 2,000 *x g* for 5 min at 4°C and supernatant retained as the substrate for PMCA. Cell lysate substrate was aliquoted and stored at -70 °C until use.

Relative amounts of PrP^C^ in DMSO and NGI-1 treated cell lysate substrate were determined by densitometric analysis of Western blot and volumes of cell lysate substrate for PMCA reactions were calculated to account for relative amounts of PrP^C^. Samples were made of cell lysate substrate (volume normalized to PrP^C^) plus 1 µL RML seed from cell lysate (above) and were brought to 100 μL total volume with PMCA conversion buffer. Master mix was made to minimize variability due to pipetting. Unsonicated samples were immediately frozen at -80 °C. Sonicated samples underwent sonication with 20 sec pulses every 30 min for 24 hr at 37 °C. Replicate PMCA reactions were done in separate sonicators.

### Immunopurification of PrP^C^ from brain tissue

3 g of CD-1 mouse brains (Biochemed, Winchester, VA, USA) were thawed and homogenized on ice in 20 mL Buffer A (20 mM MOPS pH 7.0, 150 mM NaCl) with cOmplete Protease Inhibitor Cocktail tablets (Roche) using an electric potter homogenizer. The resulting homogenate was centrifuged at 3200 *x g* for 30 min at 4 °C and the supernatant was discarded. The pellets were resuspended to a volume of 20 mL by Dounce homogenizing in Buffer A, 1% (w/v) sodium deoxycholate, 1% (v/v) Triton X-100. The homogenate was incubated on ice for 30 min to solubilize PrP^C^, then centrifuged at 100,000 *x g* for 40 min at 4 °C.

All chromatographic steps were performed at 4 °C. The solubilized supernatant was 0.45 µM filtered and passed over a column packed with 1 mL of Protein A Agarose resin (Pierce) crosslinked to D18 mAb that was pre-equilibrated with Buffer A, 1% (w/v) sodium deoxycholate, 1% (v/v) Triton X-100. The column was washed with Buffer A, 1% (w/v) sodium deoxycholate, 1% (v/v) Triton X-100. The column was eluted using 0.1 M glycine pH 2.5, 100 mM NaCl, 1% (w/v) sodium deoxycholate, 1% (v/v) Triton X-100) and pH was immediately neutralized by adding 6% (v/v) 1 M Tris pH 9.0, 5% Triton X-100, 1.4 M NaCl buffer to the eluted fractions. Fractions were dialyzed overnight against 20 mM MOPS pH 7.5, 150 mM NaCl, 0.5% (v/v) Triton X-100.

### Immunopurification of PrP^C^ from cell lysate

moRK13 cells grown in complete moRK13 media containing 1 μg/mL doxycycline were treated with 0.5 µL/mL DMSO in the media for a total of 96 hr with a media refresh at 48 hr, harvested, and processed for use as cell lysate substrate as described above (see Cell Lysate PMCA section). Cell substrate aliquots that had been stored frozen at -80 °C were thawed on ice, then subjected to an additional clarification spin at 100,000 *x g* for 40 min at 4 °C. Clarified cell supernatant was 0.45 µM filtered and passed over a 1 mL D18 mAb affinity column. All chromatographic purification steps were performed as described in “Immunopurification of PrP^C^ from brain tissue” section.

### Detection of PrP^Sc^ in PMCA reactions

Formation of PrP^Sc^ was monitored by digestion of PMCA samples with Proteinase K (PK) (Roche) and Western blotting. Samples were digested with 67 μg/mL PK at 37 °C with shaking at 750 r.p.m for 1 hr. Digestion reactions were quenched by adding SDS-PAGE loading buffer and samples were heated at 95 °C for 15 min. SDS-PAGE and Western blotting were performed as described above.

### siRNA knockdown

All siRNA experiments utilized *Silencer* Select siRNAs (ThermoFisher) and were performed using undifferentiated CAD5s in 6-well tissue culture plates. Cells were seeded to achieve 60% confluence for transfection and were allowed to adhere for a minimum of 24 hr before transfection. All knockdowns were performed using a combination of three individual targeting siRNAs (ThermoFisher, Stt3a: Cat # s68500-s68502, Stt3b: Cat # s86524-s86526). Nontargeting (scrambled) control siRNA (ThermoFisher Cat # 4390843) was used at the same final molar concentration as the combined targeting guides. For each well, 9 μL of Lipofectamine RNAiMAX transfection reagent (ThermoFisher) was diluted in 150 μL Opti-MEM. In a separate tube, 1.5 μL of each 10 μM siRNA stock (15 pmol per siRNA, 45 pmol total) was added to 150 μL of Opti-MEM. The diluted siRNA was then added to the diluted transfection reagent at a 1:1 ratio and allowed to incubate for 5 min at room temperature. 250 μL of the resulting siRNA-lipid complex was then added to each well for a total of 37.5 pmol siRNA + 7.5uL transfection reagent per well. Cells were incubated for 48 hr at 37 °C, 5% CO_2_. Uninfected CAD5 cells were assayed for surface PrP^C^ levels via flow cytometry 48 hr posttransfection. Transfection efficiency in uninfected cells was estimated via flow cytometry using a *Silencer* Cy 3-labeled GAPDH siRNA positive control (ThermoFisher Cat #AM4649) to determine effective concentration of siRNAs. Prion-infected CAD5 cells were split at a 1:5 ratio 48 hr post-transfection and allowed to adhere for 24 hr complete media. After 24 hr, prion infected cells were subjected to re-transfection using the same methodology as above and were allowed to incubate for 72 hr at 37 °C, 5% CO_2_, after which cells were lysed, processed, and analyzed for PK-resistant PrP^Sc^ via Western blot as described above.

### High speed centrifugation of CAD5 cell lysate

WT CAD5 cells were grown in triplicate in 100 mm-diameter tissue culture plates and were treated with 5 μM NGI-1, 5 μM kifunensine, or vehicle equivalent for 72 hr. After treatment, cells were washed twice with PBS and lysed in 500 μL cell lysis buffer. Protein concentrations were determined using a Pierce BCA Protein Assay Kit (Thermo Fisher Scientific) and equal amounts of total protein for each sample were ultracentrifuged at 100,000 x *g* for 1 h at 4°C in a Sorvall Discovery M120 SE Micro-Ultracentrifuge with an S45-A rotor (Thermo Fisher Scientific). The entire volume of supernatant was carefully collected. Cell pellets were resuspended in an equal volume of PBS to that of the supernatant and a Western blot for total PrP was performed using the methods described above with equal volume inputs across lanes.

### moRK13-22L treatment with NGI-1

moRK13 cells chronically infected with 22L were cultured to full confluence on 6-well plates (Corning). Replicate wells received complete growth media containing 1 μg/mL doxycycline (except for no doxycycline controls) and drug at indicated concentrations every 24-48 hrs. At experiment end point, cells were rinsed 2X with PBS and lysed as described above. PrP^Sc^ levels were determined via Western Blot as described above.

### Production of CAD5 PrP 224-Ala-MYC cells

pLNCX 224AlaMYC plasmid was kindly provided by Robert Goold (University College of London, London, UK). Plasmid-containing filter paper was soaked 100 μL TE for 30 min and 10 μL solution was added to one tube of One Shot MAX Efficiency DH5α-T1 competent cells (ThermoFisher) for transformation following manufacturer’s protocol. Cells were incubated on ice for 30 min, heat shocked at 42 °C for 30 s, then incubated on ice for 5 min. Entirety of tube was added to 2 mL S.O.C. medium (Invitrogen) and incubated at 37°C, shaking, for 1 hr. 50 μL was plated on LB agar ampicillin plates (100 μg/mL ampicillin) and incubated overnight at 37°C. The next day, one colony was selected from plate and added to 1 mL LB broth (containing 100 μg/mL ampicillin) and incubated at 37 °C, shaking at 300rpm, for 1 hr. 100 μL was added to 200 mL LB-amp broth and incubated overnight at 37°C, shaking at 300rpm. Resulting culture was maxi-prepped using Qiagen Endo-free Maxi Kit (Qiagen) according to manufacturer’s protocol. DNA concentration was estimated by NanoDrop (ThermoFisher).

CAD5 PrP KO cells were grown to 90% confluence in 6-well tissue culture plates (Corning). 4 μg pLNCX 224AlaMYC plasmid DNA was diluted in 250 μL Opti-MEM (without serum) and mixed gently. Lipofectamine 2000 was mixed gently, then 10 μL was diluted into 250 μL OptiMEM (without serum) in separate tubes. Lipofectamine was allowed to incubate for 5 min at room temperature. After incubation, diluted plasmid DNA and lipofectamine solution were gently mixed and incubated an additional 20 min at room temperature. 250 μL of the mixture was added dropwise to each well containing cells and complete medium and plate was tilted gently to mix. Cells were incubated at 37°C, 5% CO_2_ for 48 hr, after which media was changed to selection media (complete CAD5 cell media + 1 mg/mL G418). Cells were maintained in selection media for one month to establish stable lines. Monoclonal lines of the CAD5-PrP-MYC polyclonal cells were isolated via serial dilution and assayed for surface PrP^C^ expression using flow cytometry. PrP staining was done with Myc-Tag (9B11) Mouse mAb (Alexa Fluor 488 Conjugate) at 1:50 dilution (Cell Signaling Technologies). Polyclonal and monoclonal lines were frozen down in complete media containing 15% DMSO and stored in liquid nitrogen.

### Confocal microscopy for PrP^C^ and Gm1 colocalization

CAD5-PrP-MYC monoclonal cells were grown to confluence, then split 1:4 into plates containing complete media. After overnight incubation, medium was changed to complete media containing either 5 μM NGI-1 or DMSO equivalent (volume/volume). Cells were grown in drug-containing media for 72 hr, then seeded at 50,000 cells per well in drug-containing media on 6well glass-bottomed plates (No. 1.5 coverslip, 10 mm glass diameter, MatTek Cat #P06G-1.514-F) (MatTek, Ashland, MA, USA) that were coated with fibronectin. Cells were incubated at 37°C, 5% CO_2_ for 6 hr, then media was aspirated, cells were washed gently twice with PBS, and the cells that were adhered to glass were fixed in 4% paraformaldehyde in PBS (Electron Microscopy Services, Hatfield, PA, USA) for 12 min at room temperature. Following fixation, cells were washed twice with PBS and blocked for 16 hr in Fish Serum Blocking Buffer (ThermoFisher). Staining took place in PBS containing 25% Fish Serum Blocking Buffer and 0.1% Tween-20. PrP staining was done with Myc-Tag (9B11) Mouse mAb (Alexa Fluor 488 Conjugate) at 1:50 dilution (Cell Signaling Technologies) and lipid rafts were stained using Invitrogen Cholera Toxin Subunit B (Recombinant), Alexa Fluor 647 Conjugate (Invitrogen) at a 1:10,000 dilution for 1 hr at room temperature, covered from light. After antibody incubation, wells were washed with PBS and mounted using ProLong Diamond Antifade Mountant (Invitrogen). Plates were stored at room temperature, covered from light, until use.

Confocal images were acquired on a ZEISS LSM 880 scan head (ZEISS, Oberkochen, Germany) mounted on an AxioObserver stand (ZEISS), using a 1.4 NA 63X oil immersion Plan-Apochromat objective (ZEISS) and tunable filters. PrP was imaged via AF488 label with an emission window set from 495-550nm and 488nm laser excitation, cholera toxin subunit B (CTB) as a marker of lipid rafts was imaged simultaneously via AF 647 label with an emission window set from 650-735nm and 633nm laser excitation. Autofluorescence was imaged in a separate track with an emission window set from 415-465nm and 405nm laser excitation. Acquisition was 8X averaged.

Staining was analyzed in FIJI[57]. A short confocal stack was acquired and the slice closest to the coverslip was selected to maximize surface area. Cell area was determined by thresholding the autofluorescence images.

Due to low PrP labeling relative to autofluorescence, intensity was not quantified, however puncta were identified by subtracting autofluorescence estimated from the 415-465 emission band. Images were median filtered to suppress noise, and autofluorescence images were subtracted pixelwise from PrP images to highlight puncta. Puncta were not apparent in unstained controls. PrP images were binarized with a threshold at 4X the median pixel value within the cell and puncta were identified with the Analyze particles tool, excluding particles less than 5 pixels and with circularity <0.1. CTB-positive regions were similarly identified, but the autofluorescence subtraction step was omitted. PrP puncta were then assessed for overlap with CTB-positive regions.

### iPSC and cerebral organoid culture

Human-induced pluripotent stem cells (hu-iPSCs) used in this study were grown and maintained as previously described [58, 59]. In brief, hu-iPSCs (129M/V; ATCC) were cultured on low growth factor Matrigel in mTeSR1 Plus medium with 5% CO_2_ in a humidified incubator and passaged before colonies started to contact each other.

Unguided brain organoids were generated from the above iPSCs using the cerebral organoid differentiation kit (Stem Cell Technologies), based on the original protocol developed by Lancaster and Knoblich[60]. After differentiation, cultures were maintained in conical flasks on an orbital shaker at 70 rpm in complete maintenance medium: 1X GlutaMAX, 1X penicillin/streptomycin solution, 0.5% vol/vol N2, 1% vol/vol B27 with retinoic acid and 0.5X nonessential amino acids, 0.025% vol/vol insulin, and 0.00035% vol/vol 2-Merceptoethanol in 1:1 Neurobasal:DMEM-F12 medium, under standard incubator conditions (5% CO_2_, 37 °C, humidified).

### Prion infections of human cerebral organoids

Cerebral organoids were cultured for 5 months following neural induction prior to inoculation to allow for the development and maturation of astrocytes and neurons[61]. Organoid infections were performed as described previously[62]. Briefly, the culture media was replaced with fresh media containing 0.1% w/v brain homogenates from donors with type 2 (PrP-129M/V) sCJD. Unaffected brain tissue (“normal brain homogenate”, NBH) was used as a control (obtained from the NIH Neurobiobank at the University of Maryland, Baltimore, MD.). After 24 h the inoculation media was diluted 1:1 with fresh media and incubated for 6 more days. Following inoculation, organoids were transferred to new flasks containing fresh media. At either 60 or 90 dpi, organoids were treated with 10 µM NGI-1 or 0.1% DMSO as a vehicle control. Treatment containing media was changed twice a week and maintained throughout the remainder of the experiment. Organoids were collected at 120dpi and homogenized to 10% w/v in 1X PBS. Samples were cleared of cell debris via centrifugation at 2,000 x g for 2 min and stored at -80 °C.

### RT-QuIC analysis

Real-time quaking-induced conversion (RT-QuIC) assays were performed similarly to those reported previously[58, 59]. Briefly, the RT-QuIC reaction mix contained a final concentration of 10 mM phosphate buffer (pH 7.4), 300 mM NaCl, 0.1 mg/ml truncated hamster recombinant PrP90–231 (Ha90)[63], 10 μM thioflavin T (ThT), and 1 mM ethylenediaminetetraacetic acid tetrasodium salt (EDTA). Organoid homogenates were serially diluted in 10-fold steps in 0.1% SDS/1X PBS/1X N2 solution. A volume of 49μL of reaction mix was loaded into a black 384well plate with a clear bottom (Nunc), and reaction mixtures were seeded with 1 μL of the diluted homogenate for a final reaction volume of 50 μL. The final SDS concentration in the reaction, as contributed by the seed dilution was 0.002%. Reactions were run in quadruplicate for each sample. Plates were sealed (Nalgene Nunc International sealer) and incubated in a BMG FLUOstar Omega plate reader at 50 °C for Ha90 for 50 h with cycles of 60 s of shaking (700 r.p.m., double-orbital) and 60 s of rest throughout the incubation. ThT fluorescence measurements (excitation, 450 ± 10 nm; emission, 480 ± 10 nm [bottom read]) were taken every ∼45 min. Reactions were considered positive if the ThT fluorescence was greater than 10% of the maximum ThT fluorescence value on the reaction plate by the 50 h time cut-off. Spearman-Kärber analyses was used to provide estimates of the concentrations of seeding activity units giving positive reactions in 50% of replicate reactions, i.e., the 50% “seeding doses” or SD50’s as previously described[58, 64].

## Conflicts of Interest

K.S.B. and S.S. are co-inventors on a patent application related to the work in this manuscript.

## Acknowledgements

This study was funded by the National Institute for Neurological Diseases and Stroke (1R37NS125431 to SS; R01NS117276 to SS; R01NS118796 to SS) and the National Institutes of Health (P20-GM113132 to Dean Madden; T32AI007519 to KSB and Deborah Hogan). The funders had no role in study design, data collection and analysis, decision to publish, or preparation of the manuscript. We thank Dr. Joseph Contessa and Dr. Joel Watts for advice. This research was supported in part by the Intramural Research Program of the National Institutes of Health (NIH). The contributions of the NIH authors were made as part of their official duties as NIH federal employees, are in compliance with agency policy requirements, and are considered Works of the United States Government. However, the findings and conclusions presented in this paper are those of the author(s) and do not necessarily reflect the views of the NIH or the U.S. Department of Health and Human Services.

## Results

## Differences in PrP^Sc^ inhibition by NGI-1 and kifunensine are unexplained by PrP^C^ surface expression

We recently reported a genome-wide screen for regulators of PrP^C^ surface expression in the mouse neuronal cell line, CAD5, and identified N-linked glycosylation pathway genes as top hits (*manuscript in review*). Intrigued by the impact of N-linked glycosylation inhibitors on surface PrP^C^ expression, we decided to further investigate using a compound, NGI-1 [65], which inhibits the oligosaccharyltransferase (OST) complex and is further upstream in the N-linked glycan biosynthetic pathway than kifunensine, a compound previously shown to have prion-strain and cell-type dependent effects on PrP^Sc^[8] **(Fig 1A)**.

**Fig 1.**
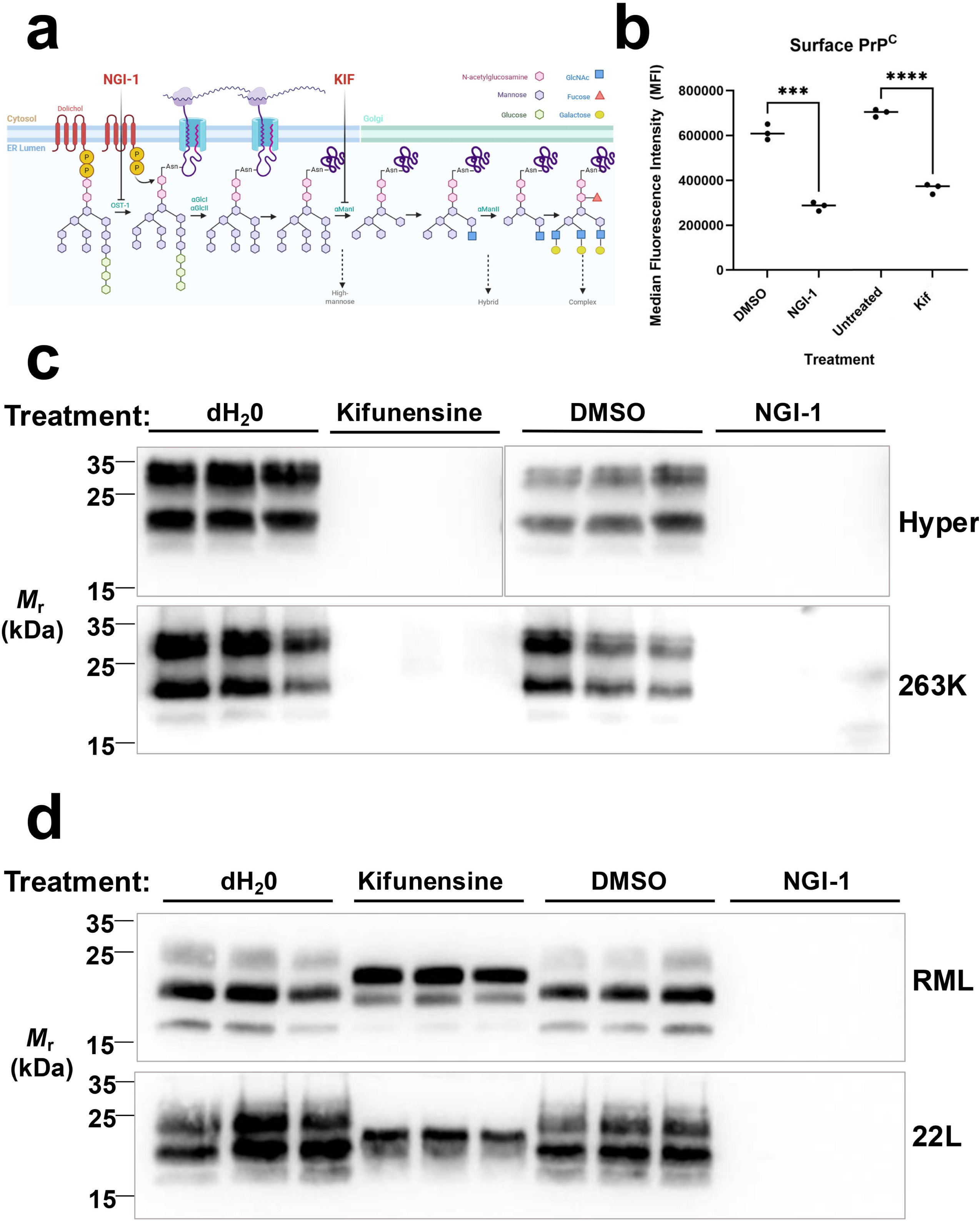
Effects of N-glycan inhibitors NGI-1 and kifunensine on PrPC surface expression in several cell types and PrPSc load in CAD5 cells chronically infected with mouse prions or hamster prions. (A) Schematic showing N-linked glycan biosynthesis in the ER/Golgi indicating enzymatic processes inhibited by NGI-1 and kifunensine. The lipid-linked oligosaccharide is transferred *en bloc* to asparagine side chains of nascent polypeptides by the enzyme oligosaccharyltransferase (OST). After transfer to the polypeptide, the glycan undergoes a series of glucose and mannose trimming steps before addition of other residues. Inhibition of discrete steps in this process is enabled using indicated small molecules, NGI-1 and kifunensine. *Beauchemin, K. (2025). Adapted from “Protein Glycosylation in the ER.” Retrieved from* https://app.biorender.com/biorender-templates (B) Cell surface PrP^C^ levels after treatment with 5 μM NGI-1 or 5 μM kifunensine in culture media for 72 hours, as measured by flow cytometry compared to vehicle-treated controls. Asterisks represent significance values from unpaired t-tests as follows: ***p 0.001, ****p 0.0001. (C) Western blot showing the loss of PK-resistant PrP^Sc^ in the lysates from CAD5 cells chronically infected with hamster Hyper or 263K prions. (D) Western blot showing the loss of PK-resistant PrP^Sc^ in the lysates from CAD5 cells chronically infected with mouse RML or 22L prions. Cells were treated with 10 μM kifunensine, 5 μM NGI-1, or vehicle controls in the cell culture media for one week. Biological triplicates are shown for each prion strain and treatment condition.

We first measured the effect of NGI-1 on PrP^C^ surface expression using flow cytometry. Addition of 5 μM NGI-1 to CAD5 cell culture media for 72 hr resulted in approximately a 50% reduction in surface PrP^C^ levels compared to vehicle-treated controls **(Fig 1B, S1A Fig)**. For comparison, we measured the effect of another N-glycosylation pathway inhibitor, kifunensine, **(Fig 1A)** on PrP^C^ surface expression. Treatment with kifunensine resulted in a similar percentage reduction in CAD5 cell surface PrP^C^ **(Fig 1B, S 1A Fig)**.

We next tested the impact of NGI-1 on PrP^Sc^ levels in CAD5 cells infected with different prion strains. Undifferentiated CAD5s expressing hamster PrP that were chronically infected with hamster prion strain Hyper (CAD5-HaPrP-Hyper) were treated with addition of 1 μM NGI-1 or 5 μM kifunensine to the media for one week with regular splitting. Western blot analysis of PK-resistant PrP^Sc^ in the cell lysates demonstrated that treatment with either kifunensine **(Fig 1C, top blot, compare dH_2_O-treated biological triplicates in lanes 1-3 to kifunensine-treated biological triplicates in lanes 4-6)** or NGI-1 **(Fig 1C, top blot, compare DMSO-treated biological triplicates in lanes 7-9 to NGI-1-treated biological triplicates in lanes 10-12)** resulted in a dramatic decrease in PrP^Sc^ load compared to their respective vehicle-treated controls **(S2A Fig for quantification)**. Treatment of CAD5 cells chronically infected with another hamster prion strain, 263K (CAD5-HaPrP-263K) yielded similar results **(Fig 1C, bottom blot, S2A Fig for quantification)**. Next, we tested undifferentiated CAD5 cells that were chronically infected with mouse prions RML (CAD5-RML) **(Fig 1D, top blot)** or 22L (CAD5-22L) **(Fig 1D, bottom blot)** using the same treatment paradigm. Strikingly, for both mouse prion strains, NGI-1 treatment resulted in a dramatic decrease of **PrP^Sc^ (Fig 1D, top and bottom blots - compare DMSO-treated biological triplicates in lanes 7-9 to NGI-1-treated biological triplicates in lanes 10-12)** while treatment with kifunensine did not **(Fig 1D, top and bottom blots – compare dH_2_O-treated biological triplicates in lanes 1-3 to kifunensine-treated biological triplicates in lanes 4-6) (S2B Fig for quantification)**. Taken together, these results suggest that (1) unlike kifunensine, the inhibitory effect of NGI-1 does not appear to be strain-specific and (2) the effects of NGI-1 on PrP^Sc^ are not solely dependent on PrP^C^ expression level at the cell surface, since treatment with either kifunensine or NGI-1 achieved similar reduction in PrP^C^ surface expression **(Fig 1B)**, but only NGI-1 decreases the abundance of mouse PrP^Sc^ **(Fig 1D)**.

### NGI-1 does not directly inhibit PrP^Sc^ formation

Using a technique called protein misfolding cyclic amplification (PMCA)[66], we examined the direct impact of NGI-1 on prion conversion by isolating this process *in vitro*. NGI-1 or DMSO was spiked directly into PMCA reactions using mouse brain homogenate as a substrate and RML from CAD5 cell lysate as the PrP^Sc^ seed. Equivalent amplification of PK-resistant PrP^Sc^ resulted from PMCA reactions containing either DMSO or NGI-1 **(Fig 2A, compare DMSO-treated lanes 2 and 8 to NGI-1 treated lanes 4 and 10)** suggesting that NGI-1 is not a direct inhibitor of PrP^Sc^ formation.

**Fig 2.**
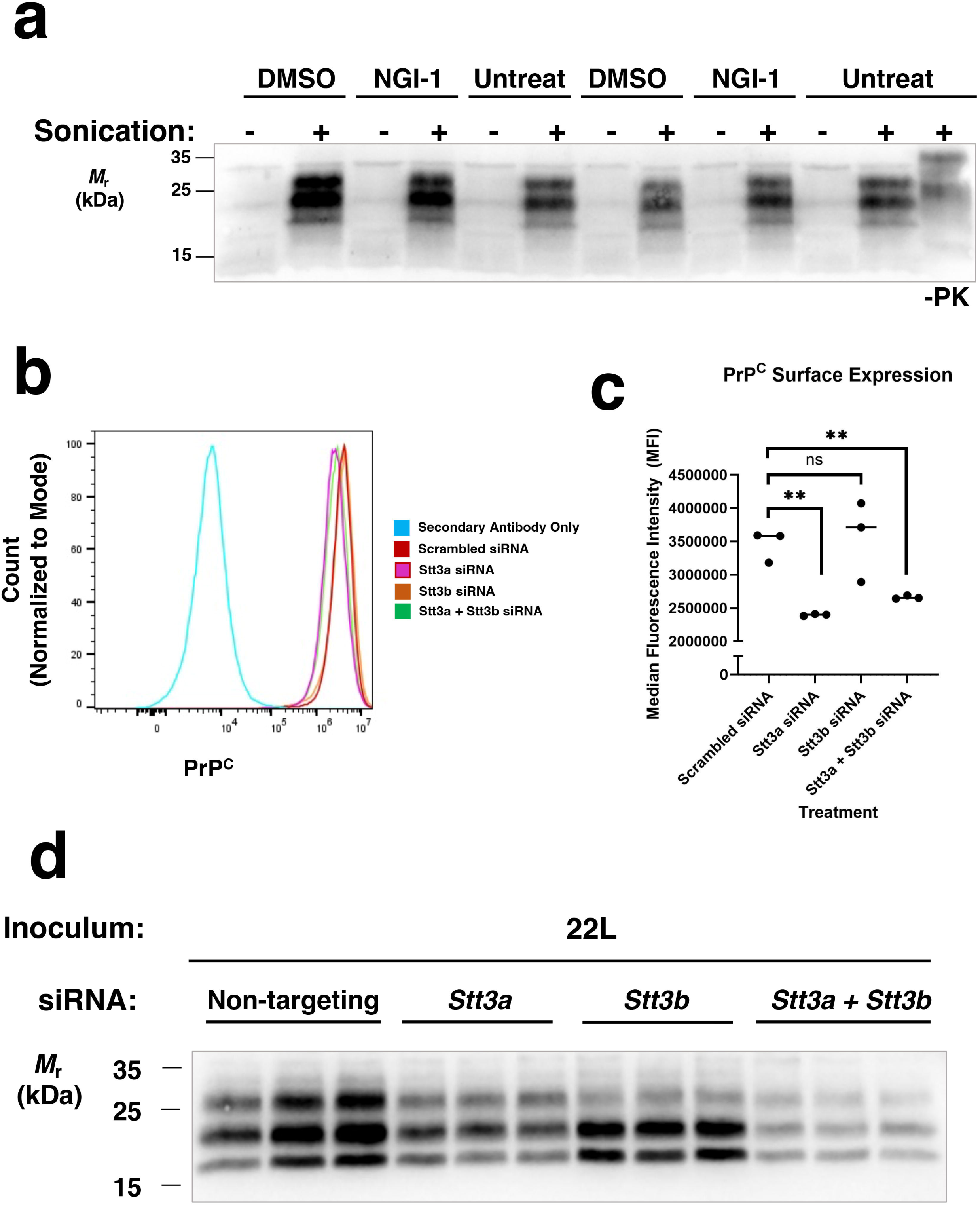
Effects of NGI-1 supplementation on PrP^Sc^ conversion in a cell-free amplification system and siRNA knockdown of targets of NGI-1 on PrP^Sc^ levels. (A) Western blot showing the amplification of PK-resistant PrP^Sc^ from brain homogenate PMCA reactions supplemented with DMSO or NGI-1. (B) Representative flow cytometry plot showing effect of siRNA knockdown of *Stt3a*, *Stt3b*, or *Stt3a* and *Stt3b* combination on surface PrP^C^ levels in WT CAD5 cells. (C) Quantification of flow cytometry PrP^C^ surface expression in undifferentiated CAD5 cells treated with siRNAs targeted at *Stt3a*, *Stt3b*, or *Stt3a* and *Stt3b* in combination. Points represent individual samples from biological triplicates. Asterisks represent significance values from unpaired t-tests as follows: **p 0.01, ns = not significant. (D) Western blot showing the loss of PK-resistant PrP^Sc^ in the lysates from CAD5 cells chronically infected with 22L prions after siRNA knockdown of *Stt3a*, *Stt3b*, or *Stt3a* and *Stt3b* in combination.

### siRNA knockdown of NGI-1’s cellular targets inhibits PrP^Sc^

NGI-1 prevents N-glycosylation by directly inhibiting two enzymes, *Stt3a* and *Stt3b*[53]. *Stt3a* and *Stt3b* are the main catalytic subunits of the OST complex (OST-A and OST-B, respectively). The OST complex is responsible for transferring a 14-sugar oligosaccharide Glc_3_Man_9_GlcNAc_2_ from dolichol to the nascent acceptor protein in the endoplasmic reticulum[47–49] **(Fig 1A)**. To determine if the inhibition of *Stt3a* and/or *Stt3b* is responsible for the inhibitory effect of NGI-1 on PrP^Sc^ levels, we utilized siRNA to knock down expression of *Stt3a*, *Stt3b*, or *Stt3a* and *Stt3b* in combination. We first looked at the effect of knockdown on PrP^C^ surface expression in uninfected CAD5 cells using flow cytometry **(Fig 2B)**. Knockdown of *Stt3a* resulted in a significant decrease in PrP^C^ surface expression **(Fig 2B and 2C, compare *Stt3a* siRNA to scrambled siRNA)**, whereas knockdown of *Stt3b* had no significant effect (**Fig 2B and 2C, compare *Stt3b* siRNA to scrambled siRNA**). Knockdown of both *Stt3a* and *Stt3b* in combination resulted in a similar decrease in PrP^C^ surface expression to knockdown of *Stt3a* alone (**Fig 2B and 2C, compare *Stt3a + Stt3b* siRNA to *Stt3a* siRNA**).

Next, we performed siRNA knockdowns of *Stt3a* and *Stt3b* in prion-infected CAD5 cells (CAD5-22L). To maintain knockdown for an extended period, siRNAs directed at either *Stt3a*, *Stt3b*, the combination of *Stt3a* and *Stt3b*, or non-targeting controls were transfected into cells twice within a week of culture. Knockdown of *Stt3a* alone resulted in a reduction of PK-resistant PrP^Sc^ by Western blot **(Fig 2D, compare biological triplicates for *Stt3a* siRNA in lanes 3-6 to biological triplicates for scrambled siRNA in lanes 1-3)**, as did knockdown of *Stt3b* alone **(Fig 2D, compare biological triplicates for *Stt3b* siRNA in lanes 7-9 to biological triplicates for scrambled siRNA in lanes 1-3)**. Interestingly, knockdown of *Stt3b* caused a shift in glycosylation pattern of the PK-resistant PrP^Sc^ with a noticeable decrease in abundance of di-glycosylated PrP^Sc^ **(Fig 2D, *Stt3b* siRNA in lanes 7-9, top band)**. Knockdown of both *Stt3a* and *Stt3b* in combination resulted in an even greater reduction in PrP^Sc^ than either *Stt3a* or *Stt3b* alone **(Fig 2D, compare biological triplicates of *Stt3a* + *Stt3b* in lanes 10-12 to biological triplicates of scrambled siRNA in lanes 1-3)**. Together, this suggests that NGI-1 is likely working to reduce PrP^Sc^ at least in part by inhibition of both *Stt3a* and *Stt3b*, and that alterations in surface PrP^C^ levels do not fully explain the observed reduction in PrP^Sc^.

### NGI-1 alters the amount and glycosylation state of PrP^C^ but does not impart large-scale changes in PrP^C^ structure or cellular localization

We wondered if NGI-1 may cause the mis-localization of PrP^C^. We confirmed that the loss of total PrP^C^ in NGI-1-treated CAD5 cell lysate is approximately 50% compared to vehicle-treated controls **(S3A Fig, S3B Fig)**. The 50% reduction in total PrP^C^ in NGI-1-treated cell lysate is consistent with the 50% reduction in surface PrP^C^ observed in NGI-1-treated CAD5 cells **(Fig 1B)**, suggesting that PrP^C^ is not retained internally because of treatment.

Next, we examined the localization of cell surface PrP^C^ in NGI-1-treated CAD5 cells. In neuronal cells, PrP^C^ is localized to areas of high cholesterol content called lipid rafts and the proper localization of PrP^C^ in lipid rafts is important for PrP^Sc^ conversion [67–71]. Thus, we decided to examine the colocalization of PrP^C^ and lipid rafts using fluorescent confocal microscopy. To directly label PrP, we utilized monoclonal cells that expressed a Myc-tagged PrP and stained with an anti-Myc-tag:AlexaFluor488 conjugate. Cholera toxin subunit B (CTB), which binds the lipid raft-associated ganglioside, GM1 [72–76], was used in a directly conjugated AlexaFluor:647 format to label lipid rafts. The degree of colocalization between PrP^C^ and lipid rafts was accessed by calculating the percentage of PrP-positive puncta within CTB-positive puncta, and no significant difference between DMSO-treated and NGI-1 treated cells was observed **(Fig 3A**, **Fig 3B)**, suggesting that NGI-1 treatment does not alter the lipid raft localization of PrP^C^ at the CAD5 cell surface.

**Fig 3.**
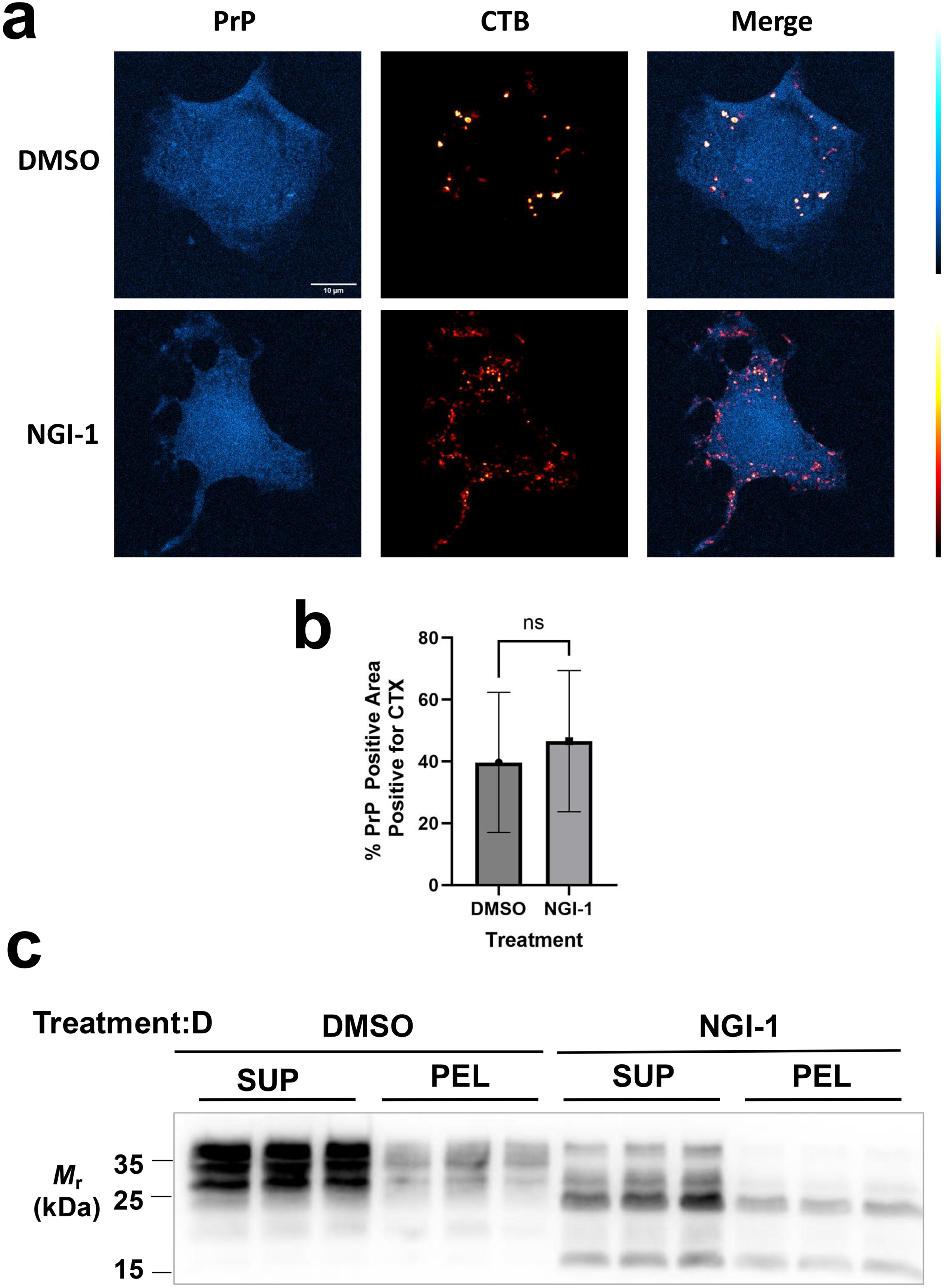
Effects of NGI-1 treatment on PrP^C^ localization and global structure. (A) CAD5-PrP-MYC cells were fixed with 4% paraformaldehyde and stained with anti-MYC antibodies to visualize PrP and cholera toxin subunit B (CTB) as a marker for lipid rafts. Single channel and merged confocal images are shown as indicated; scale bar, 10 μm. (B) Quantification of degree of colocalization between PrP puncta and CTB-positive regions. Significance values from unpaired t-tests are represented as follows: ns = not significant. (C) Western blot showing total PrP in supernatant or pellet fractions from CAD5 cells treated for 72 hr with 5 µM NGI-1 or DMSO, lysed, then subjected to centrifugation at 100,000 *x g* for 1 hr at 4°C in biological triplicate.

### NGI-1 does not significantly impact the solubility of PrP^C^

To see if NGI-1 was inducing global structural changes to PrP^C^, we performed high speed centrifugation on the lysate from CAD5 cells that had been treated with 5 μM NGI-1 or DMSO equivalent for 72 hr and examined the PrP^C^ in the resulting supernatant and pellet fractions by Western blot **(Fig 3C)**. We found the ratio of PrP^C^ found in the pellet vs. the supernatant was unchanged between treatment conditions **(quantified in S3C Fig),** suggesting that largescale structural changes to PrP^C^ (loss of soluble, globular structure) were not induced by NGI-1. Interestingly, an ∼15kDa band appeared in the NGI-1 treated cells only, which could suggest that the NGI-1-treated PrP^C^ is more susceptible to degradation by proteases. Together, these results suggest that while NGI-1 changes the total amount and glycosylation profile of PrP^C^, its anti-prion effect is not due to large scale alterations to PrP^C^ structure or localization.

### NGI-1-treated PrP^C^ does not support PrP^Sc^ amplification *in vitro*

We utilized cell lysate PMCA to determine whether NGI-1-treated cellular material could support prion conversion *in vitro*. In our first set of PMCA experiments, RK13 cells expressing mouse PrP (moRK13) under the Tet-On system were grown in the presence of doxycycline to induce PrP expression. NGI-1 or DMSO was added to the growth media for 96 hours and cell lysate prepared for use in cell lysate PMCA. Since NGI-1-treatment results in ∼50% expression of PrP^C^, we performed PMCA reactions with roughly equivalent amounts of PrP^C^ substrate **(Fig 4A, top and bottom rows, compare -PK samples in lanes 1 and 4)** and reactions with roughly twice the amount of PrP^C^ in the DMSO-treated reactions **(Fig 4A, top and bottom rows, compare -PK samples in lanes 4 and 7)**. In both cases, DMSO-treated cell lysate substrate was able to support PrP^Sc^ conversion **(Fig 4A, top and bottom rows, lanes 3 and 9)** while NGI-1-treated cell lysate was unable to support PrP^Sc^ conversion **(Fig 4A, top and bottom rows, lane 6)**.

**Fig. 4.**
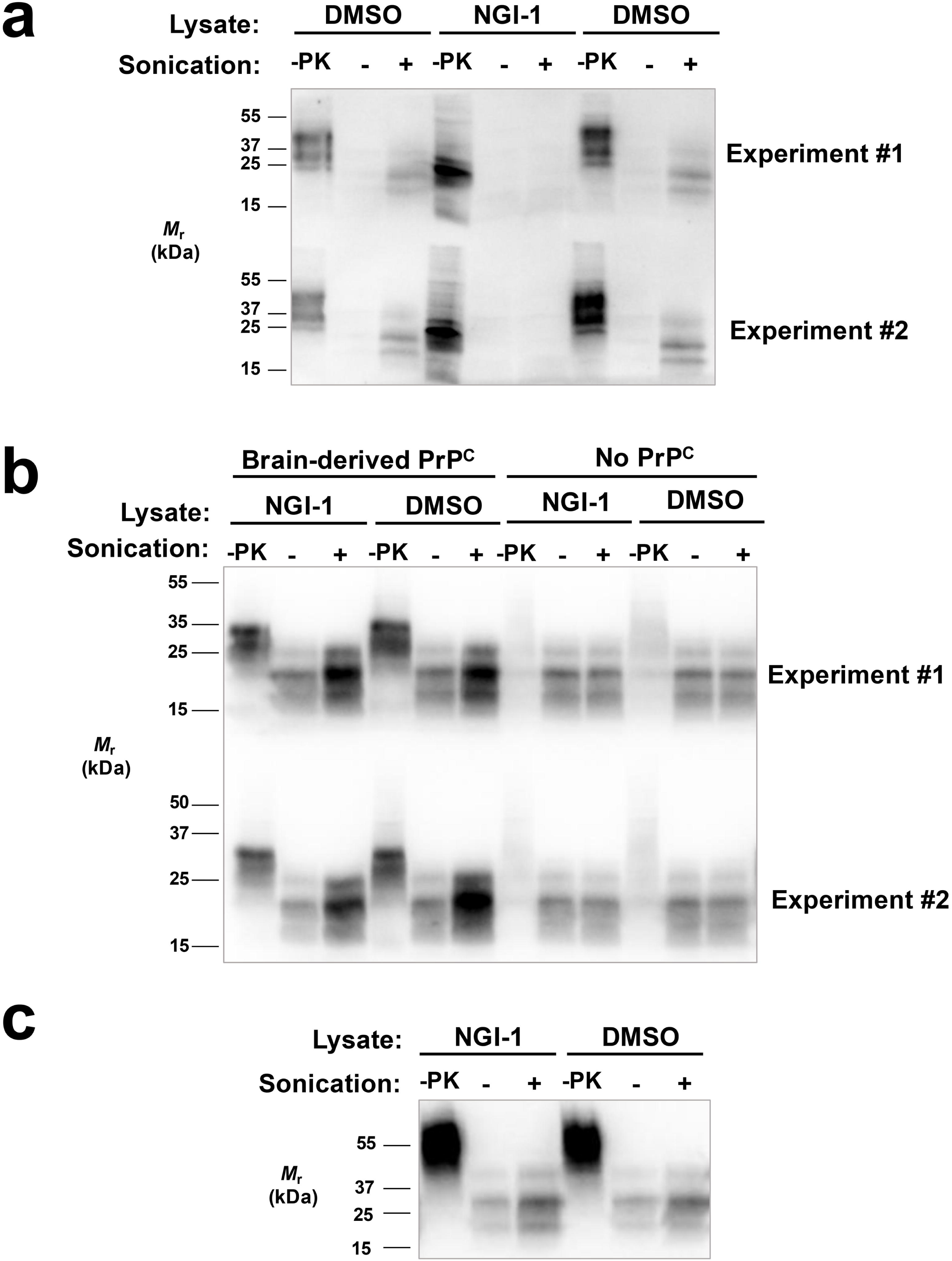
Effect of NGI-1 on amplification of PrP^Sc^ seeds in cell lysate PMCA reactions. (A) Western blot showing cell lysate PMCA reactions containing cell-derived RML seed and cell lysate substrate from DMSO or NGI-1-treated moRK13 cells expressing PrP^C^. (B) Western blot showing reconstituted PMCA reactions containing cell-derived RML seed, cell lysate substrate from DMSO or NGI-1-treated moRK13 cells that did not express PrP^C^, and immunopurified PrP^C^ from mouse brain tissue. (C) Western blot showing reconstituted PMCA reactions containing cell-derived RML seed, cell lysate substrate from DMSO or NGI-1-treated moRK13 cells that did not express PrP^C^, and immunopurified PrP^C^ from DMSO-treated moRK13 cells expressing PrP^C^.

To determine whether the inhibitory effect of NGI-1 on PrP^Sc^ conversion might be due to an indirect effect on cofactor molecules, we performed reconstituted PMCA reactions with immunopurified PrP^C^ substrate and cell lysates lacking PrP^C^. moRK13 cells were grown in the absence of doxycycline, treated with NGI-1 or DMSO as before, and treated cell lysates were used as a source of cofactor molecules in reconstituted PMCA reactions. Treated cell lysates were reconstituted with PrP^C^ purified either from mouse brain **(Fig 4B)** or moRK13 cells grown in the presence of doxycycline **(Fig 4C)**. Neither NGI-1-treated nor DMSO-treated cell lysate could support PrP^Sc^ conversion without addition of immunopurified PrP^C^ **(Fig 4B, top and bottom rows, lanes 9 and 12)**. However, both NGI-1-treated and DMSO-treated cell lysate could support PrP^Sc^ conversion when mouse brain-derived immunopurified PrP^C^ **(Fig 4B, top and bottom rows, lanes 3 and 6)** or DMSO-treated cell lysate-derived PrP^C^ **(Fig 4C, lanes 3 and 6)** was provided, suggesting that the inhibitory effect of NGI-1 is not due to an indirect effect on cofactor molecules.

### NGI-1 reduces PrP^Sc^ in non-dividing cells and is effective against human prions

We wondered whether the reduction in PrP^Sc^ observed in CAD5 cells upon NGI-1 treatment could be due to loss of PrP^Sc^ via cell division [77]. To test this, we utilized a non-dividing cell line, RK13. RK13 cells exhibit contact-dependent inhibition of division and stop dividing at full confluence. RK13 cells expressing mouse PrP (moRK13) under the Tet-On system were infected with the mouse prion strain, 22L, in the presence of doxycycline. Then, cells were split, grown to full confluence, and maintained at full confluence with addition of NGI-1 or DMSO to the growth media for one week. As with the dividing CAD5 cells, we observed a dramatic decrease in PK-resistant PrP^Sc^ via Western blot in the lysate of NGI-1-treated moRK13 cells **(Fig 5A, compare NGI-1 treated biological triplicates in lanes 13-15 to DMSO-treated biological triplicates in lanes 10-12)**. Additionally, the Western blot revealed that while DMSO-treated cells or untreated cells continued to form PrP^Sc^ throughout the treatment week **(Fig 5A, compare untreated and DMSO-treated biological triplicates in lanes 7-9 and 10-12 to untreated T=0 biological triplicates in lanes 1-3)**, NGI-1-treated cells appear to have lost PrP^Sc^ throughout the same treatment period **(Fig 5A, compare NGI-1 treated biological triplicates in lanes 13-15 to untreated T=0 biological triplicates in lanes 1-3) (S4A Fig for quantification)**.

**Fig 5.**
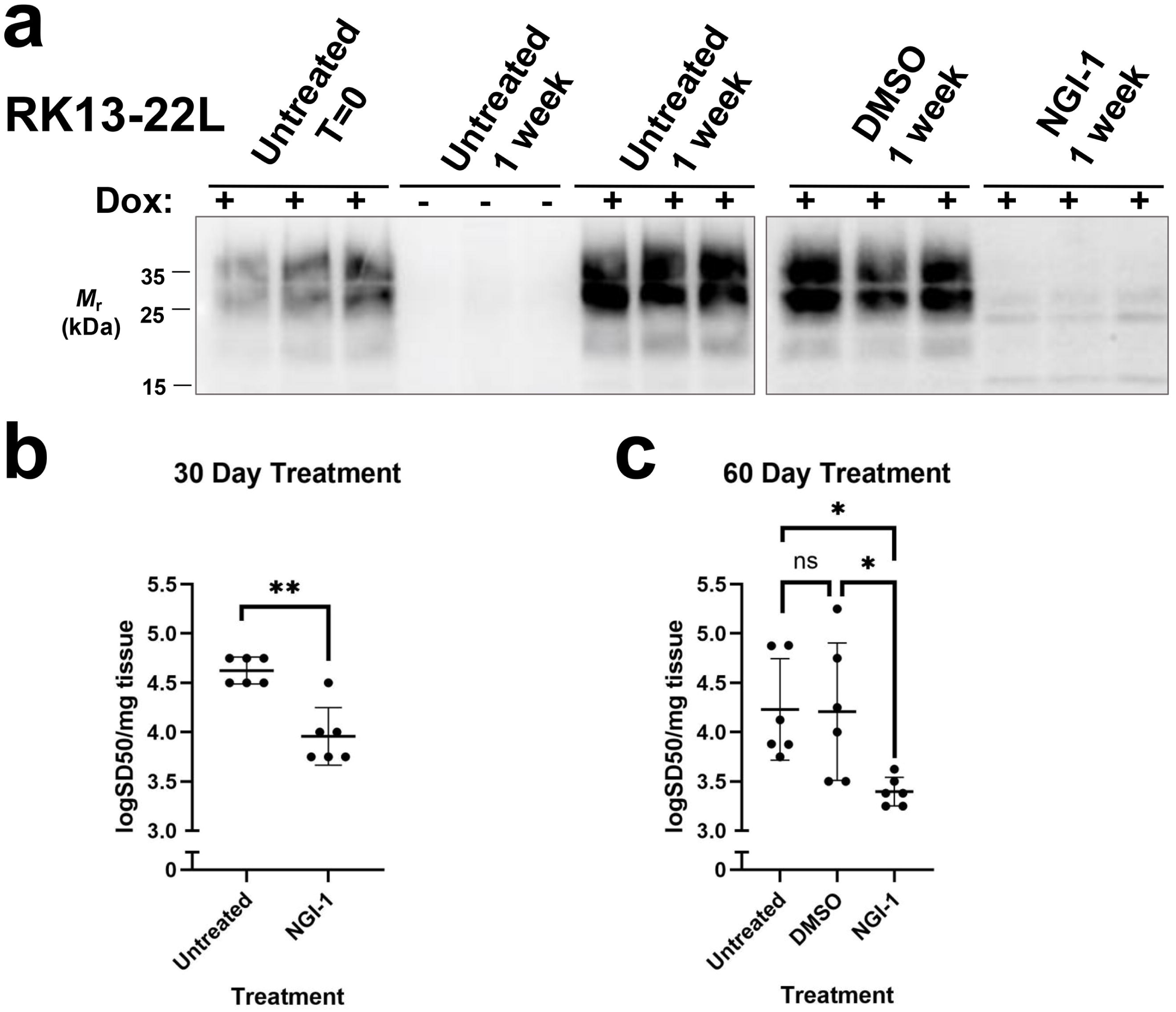
Effect of NGI-1 on PrP^Sc^ load in non-dividing cells that are chronically infected with mouse or human prions. (A) Western blot showing the loss of PK-resistant PrP^Sc^ in the lysates from non-dividing moPrP RK13 cells chronically infected with 22L prions. Cells were cultured in the presence of 1 µg/mL doxycycline for duration of experiment to induce PrP expression unless otherwise indicated. Cells were treated with 10 µM NGI-1 or DMSO equivalent for one week with no splitting. Biological triplicates are shown for each condition. (B) Log SD_50_s per mg tissue of RT-QuIC seeding activity from human sCJD-infected cerebral organoids treated therapeutically with 10 µM NGI-1 or DMSO for 30 days or (C) 60 days. Each data point represents the log SD50 from an individual organoid calculated by performing endpoint dilutions, each dilution tested in quadruplicate. Asterisks represent significance values from oneway ANOVA as follows: *p 0.05, **p 0.01, ns = not significant.

We next examined the effects of NGI-1 in human iPSCs that were fully differentiated into nondividing neurons and observed a decrease in surface PrP^C^ expression of approximately 50% **(S4B Fig, quantified in S4C Fig)**.

Human cerebral organoids have recently emerged as a valuable tool for studying sporadic Creutzfeldt Jakob disease (sCJD) prions[58, 78]. Since NGI-1 was successful in treating nondividing cells, we decided to test the effectiveness of NGI-1 as a therapeutic treatment for sCJD prions using this system. Human cerebral organoids infected with the MV2 subtype of sCJD were maintained for 60- or 90-days post infection, then treated for either 30 days **(Fig 5B)** or 60 days **(Fig 5C)** with 10 µM NGI-1 or DMSO equivalent. NGI-1 treatment caused a significant decrease in the seeding potential (logSD_50_/mg tissue) in both 30 day and 60-day therapeutic regimes **(Fig 5B and 5C)** compared to untreated or DMSO-treated controls. Taken together, these results suggest that NGI-1 can inhibit prions in non-dividing cells and can inhibit human sCJD prions.

## Discussion

We herein report a novel therapeutic target for prion disease, the oligosaccharyltransferase (OST) complex, which catalyzes the transfer of the oligosaccharide to nascent acceptor proteins during N-glycosylation[47–49]. Inhibition of the OST complex effectively depletes PrP^Sc^ in various dividing and non-dividing cell types without strain specificity. Importantly, it also effectively treats human sCJD prions in cerebral organoid models.

### Treatment with NGI-1 reduces cell surface PrP^C^ levels inhibits multiple prion strains in dividing and non-dividing cells

We observed a reduction in cell surface PrP^C^ expression of approximately 50% when mouse neuronal-like CAD5 cells were treated with NGI-1, a small molecule inhibitor of OST[65], and wondered whether a 50% reduction in cell surface PrP^C^ would translate to any noticeable reduction in PrP^Sc^ in CAD5 cells chronically infected with rodent prions. To our surprise, no detectible PK-resistant PrP^Sc^ was visible by Western blot after one week of NGI-1 treatment. Importantly, NGI-1 effectively treated CAD5s infected with various strains of mouse and hamster prions. NGI-1 treatment was also effective across various dividing and non-dividing cell types, including neurons in human organoids. That NGI-1 can effectively treat PrP^Sc^ across multiple prion strains and cell types suggests that inhibition of OST is unlikely to cause the emergence of drug-resistant prions, a challenge that has thwarted the development of many initially promising prion therapeutics[22, 23, 27–30].

Prior studies have utilized small molecule inhibitors of N-glycosylation to study PrP^C^ and PrP^Sc^[8, 29, 50], including kifunensine, swainsonine, and castanospermine, each of which inhibit enzymes further downstream in the N-glycosylation pathway than NGI-1. Every downstream inhibitor demonstrated prion strain-specific and cell-type specific effects and swainsonine treatment even caused the emergence of drug-resistant prions[29, 50].

### OST inhibition effectively treats human sCJD prions in cerebral organoids

Human cerebral organoids are infectable with human sporadic CJD (sCJD) prions[58] and have recently emerged as a viable model for sCJD drug screening[78]. We tested the OST inhibitor, NGI-1, in human cerebral organoids infected with the MV2 subtype of sporadic sCJD (sCJD) and saw significant losses in seeding capacity after 60 days or 90 days of treatment. Importantly, this demonstrates that NGI-1 treatment is an effective treatment for human sCJD prions and works against pre-established infections. These findings are significant, as many prion therapeutics have shown great promise in rodent models of disease, but were ultimately unsuccessful in treating human prions[20, 24–26, 34, 35].

Additionally, many prion diseases are not diagnosed until symptoms occur and the effectiveness of anti-prion therapeutics is highly dependent on the timing of administration[22, 25, 31–33]. Thus, the ability of NGI-1 to decrease the overall seeding capacity of sCJD-infected cerebral organoids in a therapeutic treatment paradigm (as opposed to prophylactically) gives hope that OST inhibition may be able to hinder PrP^Sc^ formation and disease progression even in symptomatic individuals. However, the degree and duration of OST inhibition required, timing of treatment initiation, and whether the loss of seeding capacity observed in the sCJD-infected cerebral organoids ultimately translates to success in treating human patients remains unknown.

### NGI-1 treatment also reduces the ability of PrP^C^ to act as a substrate for PrP^Sc^ conversion

We tested kifunensine as a control alongside NGI-1 and observed a similar 50% reduction in the cell surface expression of PrP^C^ in CAD5s treated with either drug. However, kifunensine displayed prion strain-specificity while NGI-1 did not. This key difference in strain-dependent efficacy between the two drugs suggests that the PrP^Sc^-inhibitory effect of NGI-1 cannot be solely attributable to its reduction of cell surface PrP^C^, and that there must exist a second inhibitory effect.

Our PMCA and confocal microscopy experiments indicate that the additional inhibitory effect is due to the inability of the remaining PrP^C^ molecules on the surface of NGI-1-treated cells to serve as a substrate for conversion into PrP^Sc^; rather than a direct effect of NGI-1 on PrP^Sc^ formation, NGI-1-induced changes in cofactor molecules, or mis-localization of PrP^C^ within NGI1-treated cells. The reduced ability of PrP^C^ to serve as a substrate was not accompanied by a decrease in solubility, but it is still possible that NGI-1 treatment caused a change in PrP^C^ structure that interferes with its ability to convert into PrP^Sc^.

Prior studies have shown that enzymatically deglycosylated mouse and human PrP^C^ molecules efficiently amplify PrP^Sc^ in PMCA reactions[7, 18, 19, 79]. In contrast, mice expressing unglycosylated PrP^C^ mutants have markedly reduced susceptibility to wild-type prions *in vivo*[80–83]. The reason for the discrepancy between the behavior of enzymatically deglycosylated and genetically unglycosylated PrP^C^ substrates is unknown but may be due to misfolding of unglycosylated PrP^C^ molecules within cells that does not occur when PrP^C^ molecules are deglycosylated *in vitro*. Regardless, NGI-1 treatment appears to reduce the ability of PrP^C^ molecules from forming PrP^Sc^ in PMCA reactions, cultured cells, and brain organoids.

### Combined inhibition of Stt3a and Stt3b results in synergistic anti-prion effect

To confirm that the anti-prion effects of NGI-1 treatment were due to inhibition of OST, we knocked down expression of the catalytic subunits of OST, *Stt3a* or *Stt3b,* in 22L-infected CAD5 cells using siRNA. Knockdown of *Stt3a* or *Stt3b* alone lowered PrP^Sc^ levels, while knockdown of *Stt3a* and *Stt3b* together decreased PrP^Sc^ more than knockdown of either subunit individually, suggesting that inhibition of both subunits has a synergistic effect against PrP^Sc^. Of note, NGI-1 has been shown to inhibit both *Stt3a* and *Stt3b*. We also observed that the glycoform profile of the remaining PrP^Sc^ was altered when *Stt3a* or *Stt3b* was knocked down individually. *Stt3a* knockdown caused a uniform decrease in signal across the un-, mono-, and di-glycosylated PK-resistant PrP^Sc^ bands, while *Stt3b* knockdown caused a marked decrease in the di-glycosylated band, with the un- and mono-glycosylated bands remaining relatively unchanged. This may reflect the substrate selectivity of STT3A-OST and STT3B-OST, wherein STT3A-OST is primarily responsible for glycosylation of the nascent polypeptide as it enters the lumen of the endoplasmic reticulum (ER). Following translocation, STT3B-OST glycosylates sequons skipped by STT3A, particularly carboxy-terminal glycosylation sites or those near cysteine residues[84]. Since both asparagine residues for N-glycosylation of PrP are located near the carboxyl-terminal and reside within region of the disulfide bond, it is likely that STT3B-OST plays a role in the N-glycosylation of PrP^C^. This agrees with the findings of Hirata et al., who demonstrated that PrP is imported to the ER lumen in a Sec62-dependent manner and when this pathway is utilized, STT3B-OST is typically the dominant OST isoform[85].

While prion strains and species have different characteristic glycosylation patterns and preferences[17–19, 86], glycosylation is not necessary to encode strain information[79, 87]. When PrP^Sc^ is amplified using cell free conversion methods *in vitro* with only unglycosylated PrP^C^ as a substrate, strain properties of the input PrP^Sc^ are thought to be maintained[79]. However, prior studies suggest that N-glycans have a direct impact on the kinetics of prion conversion. For example, Katorcha et al. found that sialylation slows the rate of PrP^Sc^ conversion[88]. Perhaps the loss of PrP^Sc^ upon NGI-1 treatment is due to a change in the kinetics of prion conversion, wherein the rate of formation of new PrP^Sc^ is slower than the rate of degradation of PrP^Sc^. This could partially explain the loss of PrP^Sc^ observed in non-dividing cultured cells and non-dividing human cerebral organoids.

### OST inhibitors for prion diseases: challenges and future directions

The studies herein focused on the treatment of infectious forms of prion disease. However, individuals carrying protein-altering variants in the prion protein gene, *PRNP*, may be aware of their disease risk years before onset and could potentially benefit from prophylactic therapies[89, 90]. Lowering PrP^C^ substrate is a viable therapeutic option for inherited prion disease, but how glycosylation impacts the proper folding and localization of mutated PrP^C^ in inherited prion diseases remains largely unknown[91, 92]. Interestingly, NGI-1 treatment also caused a 50% reduction in cell surface expression of PrP^C^ in fully differentiated i^3^Neurons. Thus, it is of great importance to determine whether targeting OST would be an effective treatment for inherited forms of prion disease at either pre- or post-symptomatic stages.

Unfortunately, NGI-1 is not an immediately viable candidate for anti-prion therapeutics, as it suffers from poor bioavailability[65, 93] and is unlikely to pass the blood brain barrier, making delivery a significant challenge. However, research teams are already making advances toward next-generation OST inhibitors, including some with unique specificity toward STT3A or STT3B[52, 53, 93, 94]. One could also imagine improved methods for efficient delivery of small molecules like NGI-1 to the brain or using the alternative approach of direct genetic inhibition of STT3A/B.

Human genetic data suggest that mutations in STT3A/B can lead to neurological abnormalities and intellectual disabilities[95], however it is yet unknown whether inhibition of STT3A/B in the context of a fully developed brain would mirror the effects of these mutations which were present during neuronal development and differentiation.

## Conclusions

We identified a novel therapeutic target for prion disease, the N-glycosylation enzyme complex, OST. Inhibition of OST using a small molecule, NGI-1, effectively treated PrP^Sc^ in various dividing and non-dividing cell types without prion strain-specificity. NGI-1 was also effective in treating human cerebral organoids with pre-established MV2 sCJD infection, suggesting that OST inhibition may be a viable therapeutic strategy for human prion diseases. NGI-1 appears to inhibit PrP^Sc^ formation by two synergistic mechanisms: it both lowers the abundance of cell surface PrP^C^ and reduces the ability of the remaining PrP^C^ molecules to convert into PrP^Sc^. Further development of next-generation OST inhibitors with improved pharmacokinetic properties and will be important for the future study of this novel target for anti-prion therapy.

