## Supplemental Figures for "Oligosaccharyltransferase (OST) complex inhibition effectively treats rodent and human prions"

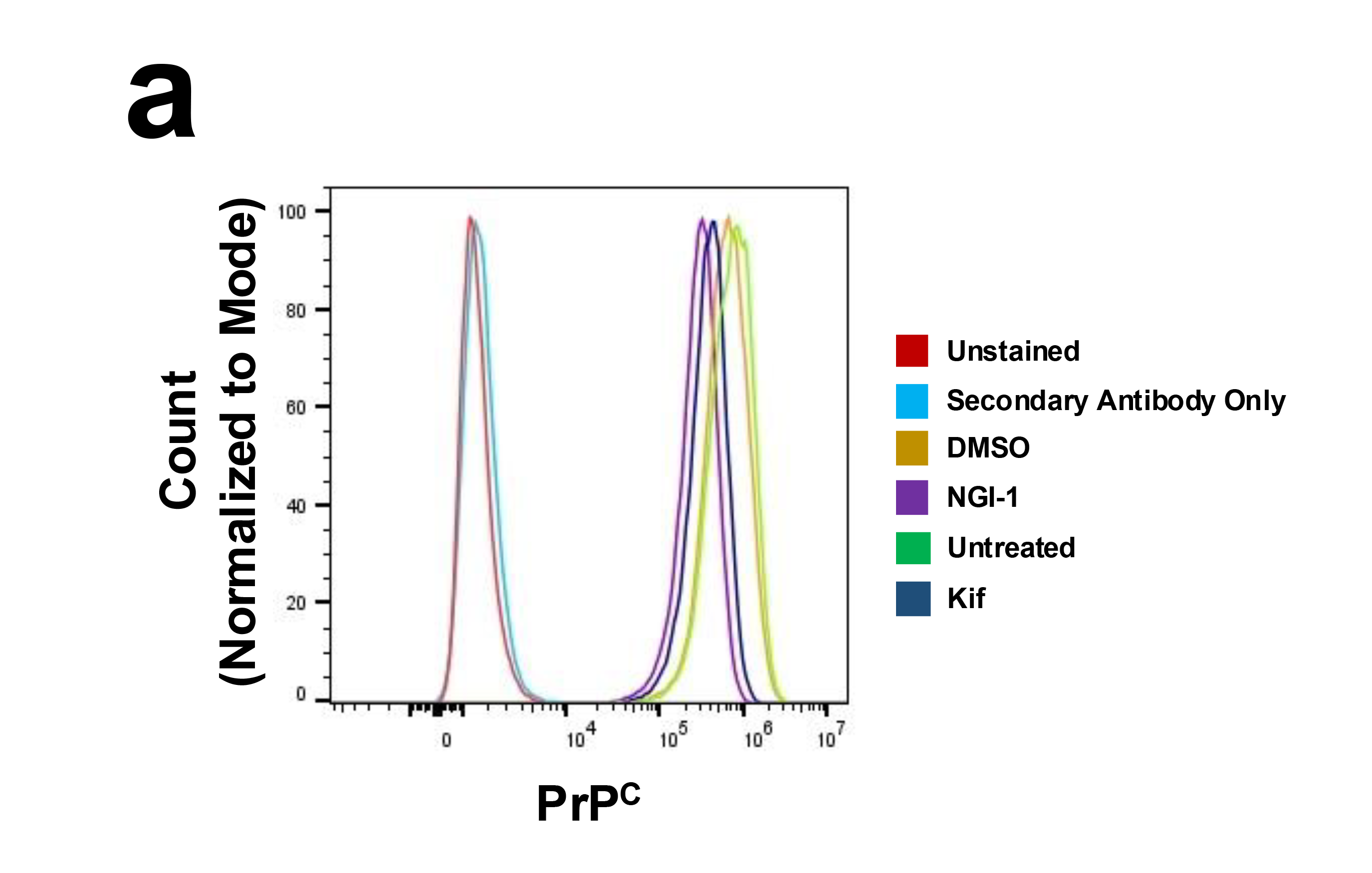

**S1 Fig. Effects of NGI-1 or kifunensine treatment on surface PrP^C^ in WT CAD5s.** (A) Representative flow cytometry plot for PrP^C^ surface expression in undifferentiated CAD5 cells treated with 5 μM NGI-1, 5 μM kifunensine, or vehicle equivalent for 72 hr.

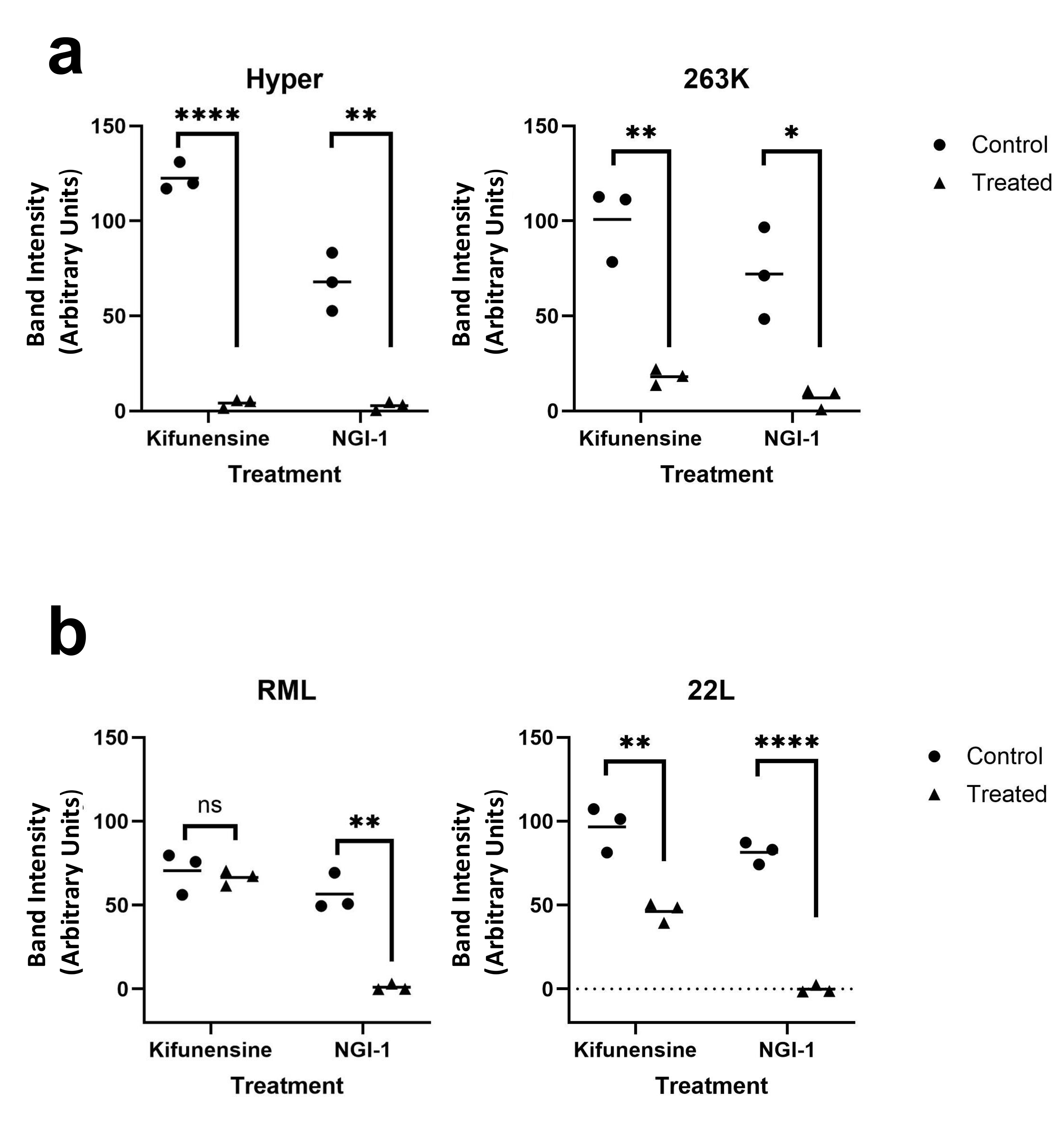

**S2 Fig. Effects of NGI-1 on PrP^Sc^ in CAD5 cells chronically infected with mouse or hamster prions.** (A) Quantification of Western blots shown in (Fig. 1C). (B) Quantification of Western blot shown in (Fig. 1D). Asterisks represent significance values from unpaired t-tests as follows: *p $\boldsymbol{\leq}$ 0.05, **p $\boldsymbol{\leq}$ 0.01, ****p $\boldsymbol{\leq}$ 0.0001, ns = not significant.

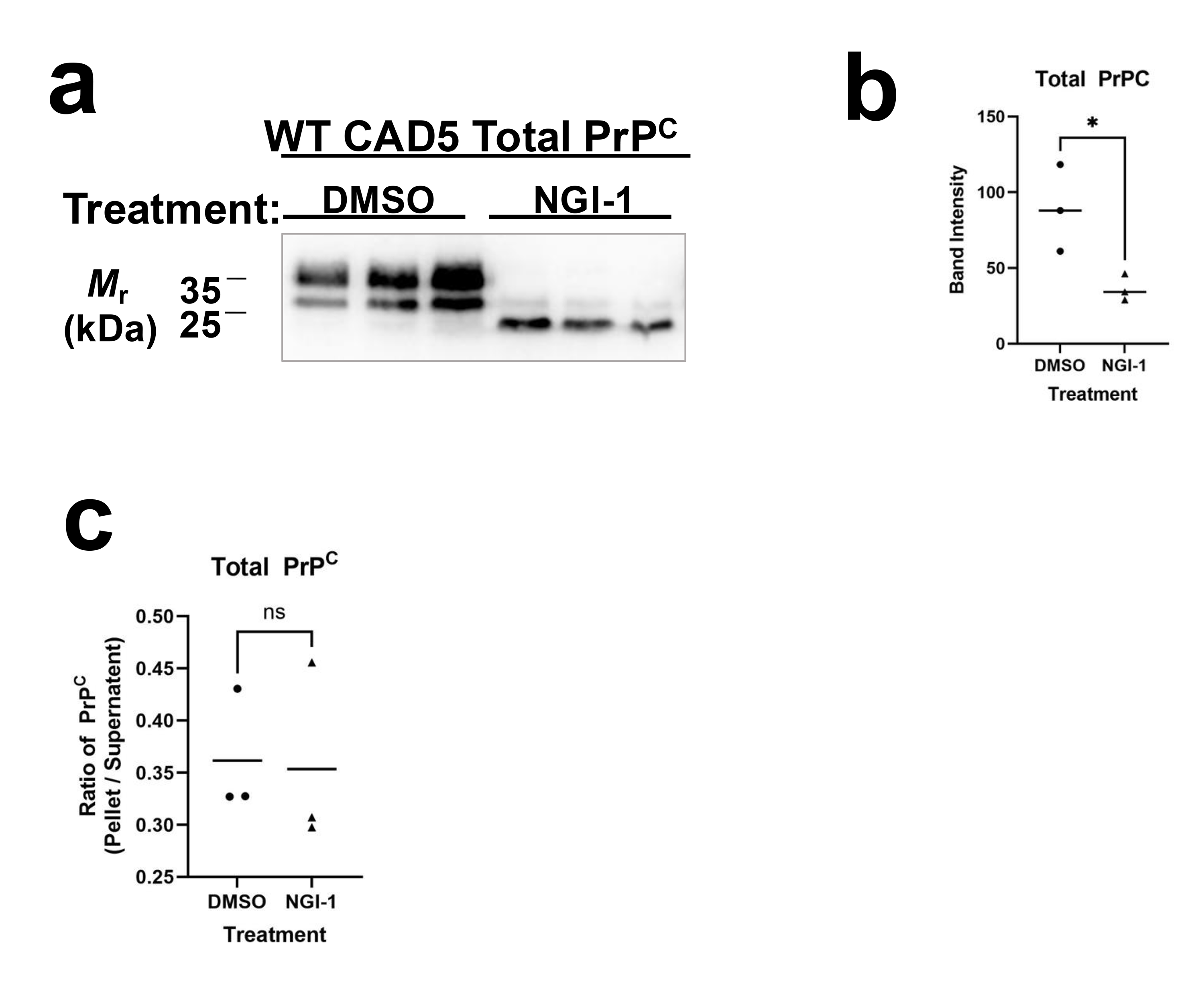

**S3 Fig. Effects of NGI-1 treatment on PrP^C^ localization and global structure.**  (A) Western blot showing total PrP in lysates from undifferentiated CAD5 cells treated for 1 week with 5 μM NGI-1 or DMSO. (B) Quantification of the Western Blot shown in (A). (C) Quantification of the Western blot in Figure 3C comparing the proportion of total PrP found in the pellet and supernatant across treatments. Points represent individual samples from biological triplicates. Asterisks represent significance values from unpaired t-tests as follows: *p $\boldsymbol{\leq}$ 0.05, ns = not significant.

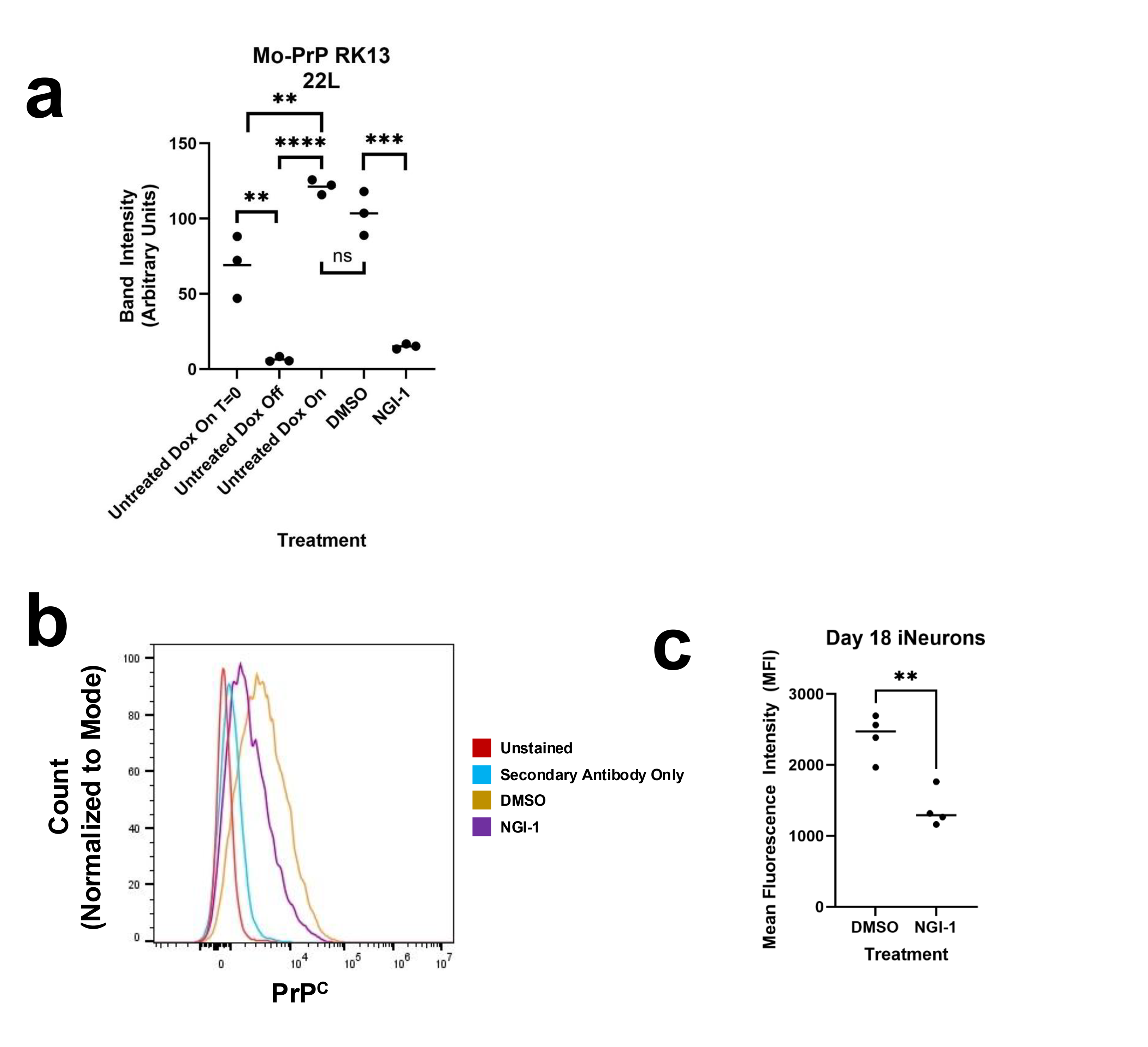

**S4 Fig. Effects of NGI-1 on PrP^C^ and PrP^Sc^ in non-dividing cells.** (A) Quantification of Western blot shown in Fig 4A. (B) Representative flow cytometry plot showing reduction of surface PrP^C^ in NGI-1-treated i^3^Neurons (C) Quantification of flow cytometry data in (B). Points represent individual samples from biological quadruplicate. Day 15 i^3^Neurons were treated with 5 μM NGI-1 or vehicle equivalent for 72 hr. Asterisks represent significance values from unpaired t-tests as follows: **p $\boldsymbol{\leq}$ 0.01, ***p $\boldsymbol{\leq}$ 0.001, ****p $\boldsymbol{\leq}$ 0.0001, ns = not significant.
